# Cooperative sensing of mitochondrial DNA by ZBP1 and cGAS promotes cardiotoxicity

**DOI:** 10.1101/2022.05.30.493783

**Authors:** Yuanjiu Lei, Jordyn J. VanPortfliet, Yi-Fan Chen, Joshua D. Bryant, Ying Li, Danielle Fails, Sylvia Torres-Odio, Katherine B. Ragan, Jingti Deng, Armaan Mohan, Bing Wang, Olivia N. Brahms, Shawn D. Yates, Michael Spencer, Carl W. Tong, Marcus W. Bosenberg, Laura Ciaccia West, Gerald S. Shadel, Timothy E. Shutt, Jason W. Upton, Pingwei Li, A. Phillip West

## Abstract

Mitochondrial DNA (mtDNA) is a potent agonist of the innate immune system; however, the exact immunostimulatory features of mtDNA and the kinetics of mtDNA detection by cytosolic nucleic acid sensors remain poorly defined. Here, we show that mitochondrial genome instability leads to Z-form mtDNA accumulation. Z-DNA Binding Protein 1 (ZBP1) stabilizes Z-form mtDNA and nucleates a cytosolic complex containing cGAS, RIPK1, and RIPK3 to sustain STAT1 phosphorylation and type I interferon (IFN-I) signaling. Increased mitochondrial Z-DNA, ZBP1 expression, and IFN-I responses are observed in cardiomyocytes after exposure to Doxorubicin, a first-line chemotherapeutic agent that induces frequent cardiotoxicity in cancer patients. Strikingly, mice lacking ZBP1 or IFN-I signaling are protected from Doxorubicin-induced cardiotoxicity. Our findings reveal ZBP1 as a cooperative partner for cGAS that sustains IFN-I responses to mitochondrial genome instability and highlight ZBP1 as a potential target in heart failure and other disorders where mtDNA stress contributes to interferon-related pathology.

## INTRODUCTION

Mitochondrial DNA (mtDNA) is increasingly recognized as an endogenous agonist of the innate immune system. Under conditions of cellular stress and mitochondrial dysfunction, mtDNA can be released into the cytoplasm or extracellular space to engage pattern recognition receptors (PRRs) including Toll-like receptor 9 (TLR9), NOD-like receptor family pyrin domain containing 3 (NLRP3), cyclic GMP-AMP (cGAMP) synthase (cGAS), and ZBP1^1–6^. The detection of mtDNA in both immune and non-immune cell types can potentiate antimicrobial innate immunity during infection but may also drive damaging inflammatory and IFN-I responses in disease and aging^7, 8^. Despite growing evidence that mtDNA is an important innate immune agonist, many questions remain. Notably, the physiological and pathological contexts of mtDNA release and the unique molecular properties of mtDNA that stabilize it in the cytosol and enhance detection by cGAS or other innate immune sensors have not been fully elucidated.

As a covalently closed, circular genome, mtDNA lacks free DNA ends that can rotate to relieve torsional strain induced during replication and transcription^9^. Consequently, the activity of mitochondrial replication and transcription machineries can generate overwound (positively supercoiled) and underwound (negatively supercoiled) regions of mtDNA, which may promote the formation of other immunostimulatory mtDNA structures if not properly resolved. Z-DNA is one such non-canonical structure and is a left-handed helix that differs from canonical Watson–Crick B- DNA^10, 11^. Sequences with alternating purine–pyrimidine regions have the highest propensity to form Z-DNA, although negative supercoiling of other sequences can promote B to Z transition^12, 13^. In mitochondria, the activity of topoisomerases DNA topoisomerase I, mitochondrial (TOP1MT) and DNA topoisomerase III alpha (TOP3A) are required to relieve torsional mtDNA stress and relax negative supercoiling^9, 11^. However, the degree to which torsional stress promotes B- to Z-DNA transition in mtDNA is unknown, and whether the acquisition of this unique DNA confirmation enhances or sustains the immunostimulatory potential of mtDNA is presently unclear. Emerging evidence suggests ZBP1 is a critical innate immune receptor for both nuclear Z-DNA and double stranded Z-RNA^14–22^, yet a role for ZBP1 in sensing Z-form mtDNA has not been explored.

Type I interferons (IFN-I) are pleotropic cytokines that function in antimicrobial immunity, tumor immunity, and metabolic regulation. However, persistent and unrestrained IFN-I signaling is linked to autoimmune diseases, type I interferonopathies, cancer, and aging-related disorders^8, 23–26^. For example, mtDNA sensing by cGAS or TLR9 has been linked to IFN-I and inflammatory responses in heart failure^8, 27–31^. Interestingly, the off-target effects of radiotherapy and chemotherapy can cause cardiomyopathy in cancer survivors^32^, which is associated with aberrant innate immune activation^33, 34^. Anthracycline chemotherapeutics such as Doxorubicin (Doxo) are the leading causes of cancer therapy-related heart failure^35–38^. The molecular and cellular mechanisms responsible remain elusive, although mitochondrial dysfunction and mtDNA damage are hypothesized to be major drivers of Doxo-induced cardiotoxicity (DIC)^39–44^. Recent work has shown that Doxo can induce mitochondrial genome instability, mtDNA supercoiling, and IFN-I in vitro^45–47^. However, a direct role for ZBP1 in cardiac IFN-I signaling and/or DIC has not been explored.

Here, we report that ZBP1 is a key innate immune sensor of mitochondrial genome instability. Induction of torsional stress in mtDNA promotes the formation of mitochondrial Z-DNA. ZBP1 stabilizes Z-form mtDNA and nucleates a cytosolic complex containing cGAS, receptor-interacting protein 1 (RIPK1), and RIPK3 to engage robust IFN-I signaling. Moreover, we show that Doxo induces ZBP1-dependent IFN-I program in cardiomyocytes and cardiac myeloid populations in vivo. Ablation of ZBP1 or IFN-I signaling limits cardiotoxicity and improves heart function during and after administration of Doxo chemotherapy, shedding new light on the immune mechanisms of DIC and revealing a potential therapeutic avenue for cancer therapy-associated heart failure.

## RESULTS

### ZBP1 is required for robust and sustained IFN-I responses to mitochondrial genome instability

To first explore roles for additional IFN-I-induced nucleic acid sensors in mtDNA sensing, we mined RNA expression data from the TFAM heterozygous knockout model (*Tfam^+/-^*), which exhibits persistent mitochondrial genome instability, cytosolic mtDNA release (Figure S1A), and sustained IFN-I signaling^48^. Analysis of *Tfam^+/-^* mouse embryonic fibroblasts (MEFs) revealed elevated expression of ZBP1, IFI200 family members, and mitochondrial antiviral signaling (MAVS)- dependent RNA helicases (Figures 1A and S1B). Consistent with our prior work documenting cytosolic mtDNA as the IFN-inducing ligand in TFAM-deficient cells, *Sting^-/-^* MEFs displayed markedly reduced ISG expression after TFAM silencing (Figure S1C). Conversely, MAVS-null MEFs maintained high levels of ISGs after TFAM silencing, indicating that mitochondrial RNA is not a major driver of IFN-I signaling in TFAM-deficient cells (Figure S1C).

Knockdown of putative DNA sensors *Ifi203*, *Ifi204*, or *Ifi205* did not reduce TFAM-induced ISG expression (data not shown); however, silencing of ZBP1 markedly attenuated ISG expression in *Tfam^+/-^* cells (Figure S1D). We next obtained ZBP1-null mice^49^ and crossed them onto the *Tfam^+/-^* background. RNA-seq analysis indicated that in contrast to MEFs from *Tfam^+/-^* mice, *Tfam^+/-^Zbp1^-/-^* cells failed to sustain elevated expression of ISGs triggered by mtDNA instability (Figure 1A and Table S1). Additional qRT-PCR and western blotting experiments confirmed that loss of ZBP1 was sufficient to largely abrogate ISG expression in *Tfam^+/-^* cells (Figures 1B and 1C). Experiments in human cells further substantiated the key role for ZBP1 in sensing mtDNA genome instability (Figure 1D). In addition, experiments in 293FT cells stably reconstituted with STING (293FT-STING) revealed that cells expressing ZBP1 alone failed to express ISGs after TFAM silencing, whereas cells co-expressing cGAS and ZBP1 showed augmented ISGs compared to cells expressing cGAS alone (Figure 1E). Collectively, these results indicate that ZBP1 is an essential mtDNA sensor that synergizes with cGAS to sustain IFN-I responses to mtDNA genome instability.

**Figure 1.**
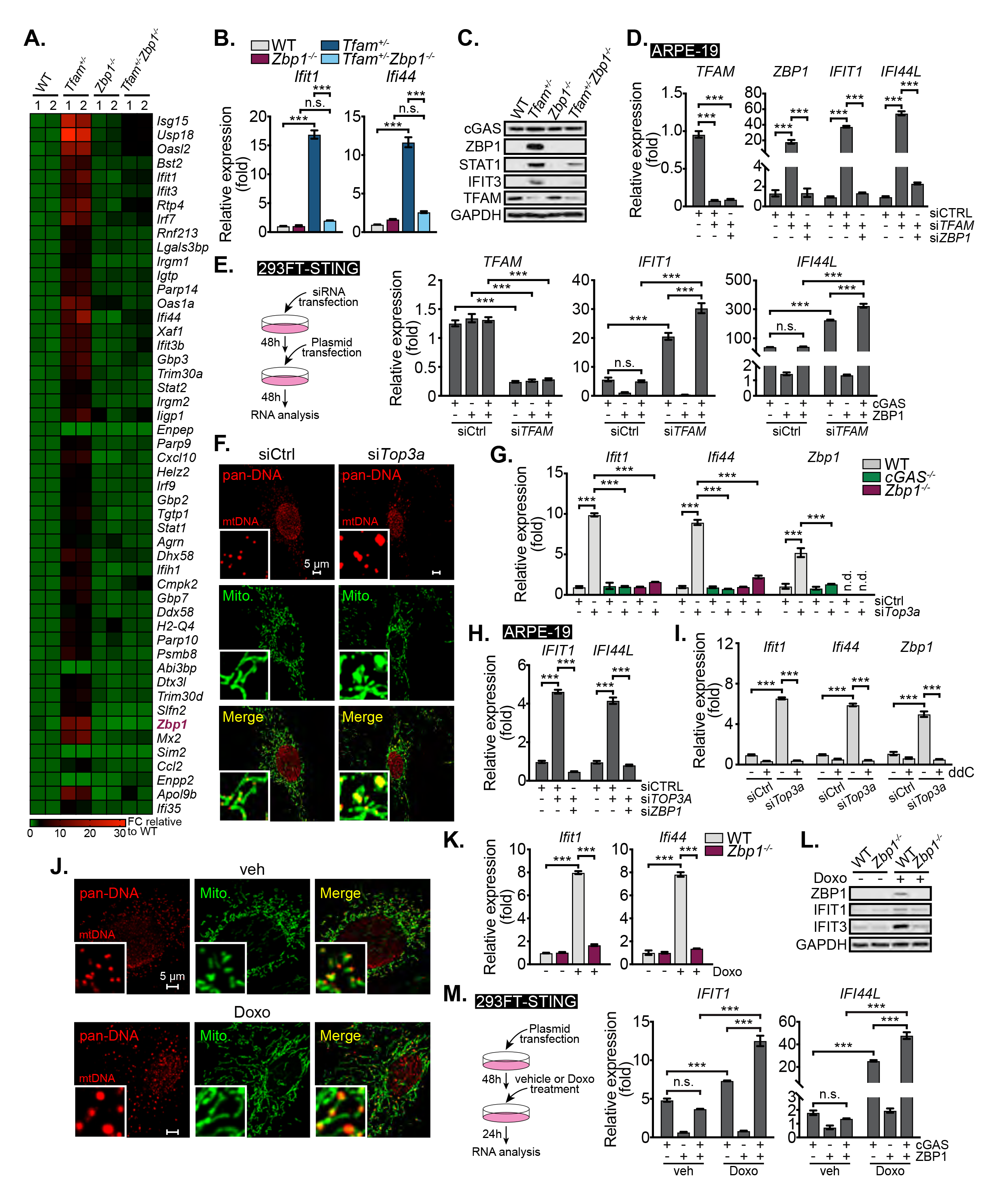
ZBP1 sustains IFN-I responses to mitochondrial genome instability. **A**, RNA-seq heatmaps showing the top 50 genes with the lowest p-values between WT and *Tfam^+/-^* genotypes. Fold changes (FC) are normalized to the average of WT MEFs. **B, C**, qRT-PCR analysis (n = 3) (**B**) and western blots (**C**) of ISGs in WT, *Tfam^+/-^*, *Zbp1^-/-^* and *Tfam^+/-^Zbp1^-/-^* MEFs. **D**, qRT-PCR analysis (n = 3) of *TFAM* and ISGs in ARPE-19 transfected with siCTRL, si*TFAM* or si*TFAM* + si*ZBP1* for 72 h. **E**, Schematic illustration of siRNA and plasmid transfection in 293FT-STING (left). qRT-PCR analysis (n = 3) of TFAM and ISGs in 293FT-STING transfected with indicated siRNA and plasmid (0.25 μg/mL cGAS, 0.25 μg/mL ZBP1 or equivalent amount of empty vector) (right). **F**, Representative microscopy images of WT MEFs transfected with siCtrl or si*Top3a* and stained with anti-pan-DNA and -HSP60 (Mito.) antibodies. Inset panels are magnified 4x. **G**, qRT-PCR analysis (n = 3) of ISGs in WT, *cGAS^-/-^* and *Zbp1^-/-^* MEFs transfected with siCtrl or si*Top3a* for 96 h. **H**, qRT-PCR analysis (n = 3) of ISGs in ARPE-19 transfected with siCTRL, si*TOP3A* or si*TOP3A* + si*ZBP1* for 72 h. **I.** qRT-PCR analysis (n = 3) of ISGs in WT MEFs transfected with siCtrl or si*Top3a* with or without 2’,3’- dideoxycytidine (ddC, 100 μM) for 72 h. **J**, Representative confocal microscopy images of WT MEFs treated with vehicle (veh) or Doxorubicin (Doxo, 500 nM, 24 h) and stained with anti-pan-DNA and - HSP60 (Mito.) antibodies. Inset panels are magnified 4x. **K, L**, qRT-PCR (n = 3) analysis (**K**) and western blots (**L**) of ISGs in WT and *Zbp1^-/-^* MEFs treated with or without Doxo (50 nM, 48 h). **M.** Schematic illustration of plasmid transfection and Doxo treatment in 293FT-STING (left). qRT-PCR analysis (n = 3) of ISGs in 293FT-STING transfected with indicated plasmids (0.4 μg/mL cGAS, 0.8 μg/mL ZBP1 or equivalent amount of empty vector) and with or without Doxo (200 nM) treatment (right). Statistical significance was determined using analysis of variance (ANOVA) and Tukey post hoc test (**A, D, E, G, H, I, L, M**). *P < 0.05, **P < 0.01, and ***P < 0.001. Error bars represent SEM. See also Figure S1 and Table S1.

Reduced TFAM expression can alter mtDNA topology, promoting the transition from predominantly relaxed monomeric circles to catenated^50^ or supercoiled mtDNA species (Figure S1E). The mitochondrial isoform of TOP3A functions to resolve mtDNA catenation and relax negative supercoiling induced during mtDNA replication and transcription^9, 11, 51, 52^. As loss of TOP3A causes mtDNA torsional strain leading to mitochondrial nucleoid aggregation and genome instability similar to TFAM depletion (Figure 1F)^51^, we next investigated whether silencing of TOP3A promotes mtDNA-dependent IFN-I signaling. Interestingly, siRNA knockdown of TOP3A in both mouse and human cells (Figures S1F and S1G) triggered robust ISG expression in a cGAS- and ZBP1-dependent fashion (Figures 1G and 1H). Selective depletion of mtDNA with the chain terminating nucleoside 2’3’-dideoxycytosine (ddC) during TOP3A knockdown revealed that mtDNA, and not nuclear DNA, is the endogenous ligand driving cGAS- and ZBP1-dependent IFN-I responses (Figures S1H and 1I). Mammalian mitochondria also house the type IB topoisomerase TOP1MT, which facilitates mtDNA decatenation by TOP3A and resolves positive and negative mtDNA supercoiling^11, 52–54^. Similar to results obtained by targeting TOP3A, siRNA knockdown of TOP1MT triggered ISG expression in a cGAS-, ZBP1-, and mtDNA-dependent fashion (Figures S1I-S1L). Collectively, these data indicate that loss of multiple mitochondrial factors governing mtDNA packaging and topology triggers ISG expression via both cGAS and ZBP1.

Doxorubicin is a widely used anthracycline chemotherapeutic drug that induces mtDNA damage and mitochondrial dysfunction, in addition to nuclear DNA damage^39, 40^. A recent study revealed that Doxo promotes mtDNA supercoiling and catenation^47^, and we observed that Doxo induced mtDNA nucleoid aggregation similar to cells lacking TFAM or TOP3A (Figures 1J). Doxo treatment also promoted cytosolic mtDNA release, as well as upregulation of ZBP1 and other ISGs (Figures S1M-S1O). Moreover, Doxo-induced ISG expression was independent of MAVS and RNA sensing but required signaling via the type I interferon receptor (IFNAR) and mtDNA (Figures S1P- S1R). Doxo-induced ISG expression was also markedly impaired in *Zbp1^-/-^* cells, similar to TFAM, TOP3A, and TOP1MT knockdown cells (Figures 1K and 1L). Pretreatment of *Zbp1^-/-^* cells with interferon beta (IFNβ) before Doxo challenge failed to increase ISGs to the same level as Doxo-treated WT cells, indicating that another IFN-I induced DNA sensor cannot compensate for the loss of ZBP1 (Figure S1S). Finally, reconstitution experiments in 293FT-STING cells revealed that cGAS and ZBP1 synergize to enhance ISG expression in response to Doxo (Figure 1M). Despite these crucial roles for ZBP1 in promoting IFN-I responses downstream of mtDNA instability, we noted no requirement for ZBP1 in sensing linear B-form immunostimulatory DNA, as reported by several other groups^14, 16, 49, 55^ (Figure S1T). Moreover, ZBP1 was not required for cGAS-dependent ISGs triggered by inner mitochondrial membrane (IMM) herniation and caspase inhibition^56–58^ (Figure S1U). In sum, these results suggest that ZBP1 is a unique sensor of mitochondrial genome instability that augments cGAS- STING dependent IFN-I signaling in human and mouse cells.

### mtDNA genome instability promotes the accumulation of Z-DNA that is stabilized by ZBP1

Negatively supercoiled plasmid DNA can form stable Z-DNA in vitro^59–61^. Moreover, topoisomerase insufficiency can promote the formation of non-canonical DNA structures, including left handed Z-DNA^11, 62^. Bidirectional replication and transcription of circular mtDNA promotes supercoiling that must be resolved by TOP1MT and TOP3A. This raises the possibility that increased negative mtDNA supercoiling in TOP3A or TOP1MT deficient cells, and perhaps *Tfam^+/-^* and Doxo-treated cells, promotes the formation of stable Z-DNA for detection by ZBP1. To explore this hypothesis, we employed a Z-DNA antibody that selectively binds Z-DNA in vitro and in cellulo^61, 63, 64^. To first validate the selective affinity of the Z22 antibody for Z-DNA, we generated Z-DNA in vitro by hybridizing two 74 bp ssDNA minicircles^61^ (Figure 2A). Gel shift and slot blotting assays revealed that the Z22 antibody strongly binds plasmid Z-DNA, but not a plasmid with the identical sequence in B- DNA confirmation (Figures 2B, S2A and S2B). Moreover, the Z22 antibody efficiently labeled plasmid Z-DNA in the cell cytoplasm after transfection, but did not immunostain B-DNA, highlighting the utility of this reagent for fluorescent microscopy analysis of Z-DNA (Figure S2C).

**Figure 2.**
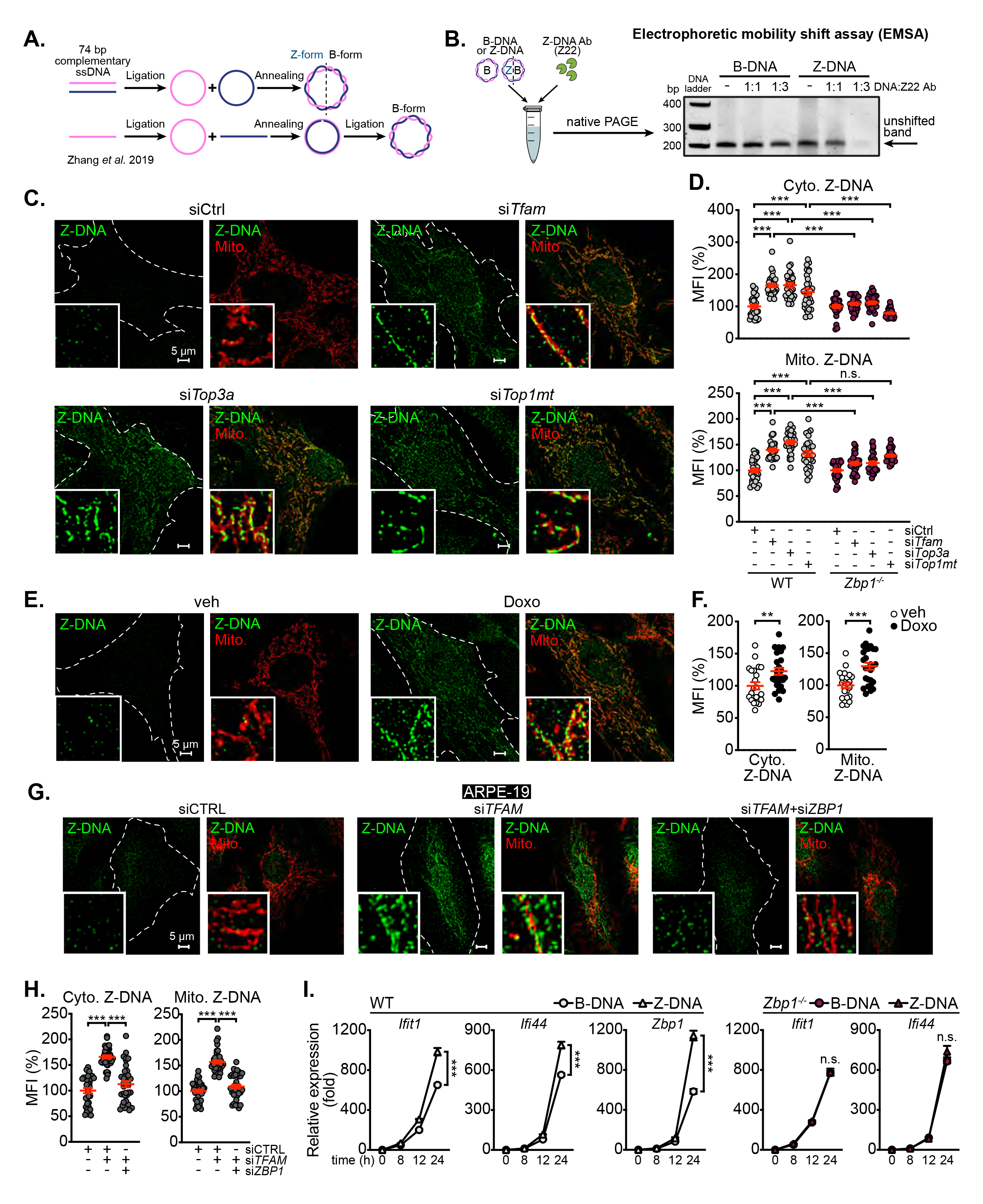
Mitochondrial genome instability promotes Z-form DNA accumulation that is stabilized by ZBP1. **A**, Schematic illustration of circular Z-form and B-form DNA synthesis. **B**, Electrophoretic mobility shift assay (EMSA) measurement of binding of Z-DNA-specific antibody Z22 to B-DNA and Z-DNA. DNA and Z22 are mixed at molar ratio of 1:1 or 1:3. **C**, Representative images of WT MEFs transfected with siCtrl, si*Tfam*, si*Top3a* or si*Top1mt* for 72 h and stained with anti-Z-DNA and -TOMM20 (Mito.) antibodies. Inset panels are magnified 4x. **D**, Mean fluorescent intensity (MFI) quantification of cytosolic (Cyto.) and mitochondrial (Mito.) Z-DNA in WT and *Zbp1^-/-^* MEFs transfected with siCtrl, si*Tfam*, si*Top3a* or si*Top1mt* for 72 h (n ≥ 30 fields for each group from n = 3 experiments). MFI percentiles are normalized to WT or *Zbp1^-/-^* MEFs transfected with siCtrl. **E, F**, Representative images of WT MEFs treated with vehicle (veh) or Doxorubicin (Doxo) and stained with anti-Z-DNA and -TOMM20 (Mito.) antibodies (**E**). MFI of Cyto. and Mito. Z-DNA is quantified in (**F**). (n ≥ 20 fields for each group from n = 3 experiments). MFI percentiles are normalized to veh-treated WT MEFs. **G, H**, Representative images of ARPE-19 transfected with siCTRL, si*TFAM* or si*TFAM* + si*ZBP1* for 72 h and stained with anti-Z-DNA and -TOMM20 antibodies (**G**). Inset panels are magnified 4x. MFI of Cyto. and Mito. Z-DNA is quantified in (**H**) (n ≥ 30 fields for each group from n = 3 experiments). MFI percentiles are normalized to ARPE-19 transfected with siCTRL. **I**, qRT-PCR analysis (n = 3) of ISGs in WT and *Zbp1^-/-^* MEFs transfected with B-DNA or Z-DNA (125 ng/mL) for indicated time. Statistical significance was determined using analysis of variance (ANOVA) (**D, I**) and Tukey post hoc test (**D**), or unpaired t-test (**F**). *P < 0.05, **P < 0.01, and ***P < 0.001. Error bars represent SEM. See also Figure S2.

Utilizing the Z22 antibody and quantitative immunofluorescence microscopy, we next screened for the presence of Z-DNA in cells experiencing mtDNA genome stress. Strikingly, we observed increased cytosolic and mitochondrial Z-DNA accumulation in TFAM, TOP3A, and TOP1MT knockdown cells (Figures 2C and 2D). Doxo also enriched Z-DNA in the cytosol and mitochondria as revealed by microscopy and slot blotting (Figures 2E, 2F and S2D), further supporting the notion that perturbations enhancing mtDNA supercoiling promote the accumulation of Z-DNA. Depletion of mtDNA with ddC after silencing of TOP3A or TOP1MT, or Doxo treatment, reduced both mtDNA nucleoid abundance and aggregation (Figure S2E), while also significantly decreasing overall Z-DNA intensity (Figure S2F), strongly indicating that mtDNA was the source of Z-DNA we observed. However, IMM herniation and caspase inhibition were insufficient in inducing cytosolic or mitochondrial Z-DNA accumulation (Figure S2G). Additional experiments in human cells confirmed increased Z-DNA abundance after TFAM silencing (Figures 2G and 2H). ZBP1 can stabilize Z-form DNA via its Z-DNA binding domains^65–68^, and therefore we next explored whether ZBP1 expression is required for robust Z-DNA accumulation in cells experiencing mtDNA stress. Interestingly, we observed less cytosolic Z-DNA after TFAM, TOP3A, or TOP1MT silencing in *Zbp1*^-/-^ MEFs relative to WT MEFs, with similar trends seen after dual silencing of TFAM and ZBP1 in human cells (Figures 2D, 2G and 2H). Finally, exposure of *Tfam^+/-^*MEFs to hydralazine, a vasodilator that is known to stabilize Z-form DNA to cause drug-induced lupus^69, 70^, dramatically increased overall Z-DNA staining, as well as ISG expression in a ZBP1-dependent fashion (Figures S2H and S2I).

We next sought to determine whether Z-DNA is intrinsically more immunostimulatory to the cGAS-IFN-I axis. Transient transfection of either plasmid Z-DNA or B-DNA into WT cells elicited comparable ISG expression at early timepoints. However, Z-DNA was significantly more stimulatory at 12-24 hours post transfection (Figure 2I). Z-DNA and B-DNA elicited similar levels of ISGs in *Zbp1^-/-^* cells, suggesting that the Z-DNA stabilizing and/or signaling functions of ZBP1 are necessary to augment IFN-I signaling. Additional studies in ARPE-19 cells confirmed findings in MEFs, revealing that transfected plasmid Z-DNA is more stimulatory than B-DNA in human cells (Figure S2J). Finally, using an in vitro cGAS activity assay^71^, we observed no differences in cGAMP production in the presence of either plasmid, indicating that cGAS enzymatic activity is comparably induced by both B- and Z-DNA (Figure S2K). Collectively, these data show that Z-DNA is a potent IFN-I inducer and reveal that torsional mtDNA stress promotes the formation of Z-DNA that is stabilized by ZBP1.

### ZBP1 and cGAS form a DNA- and RHIM-dependent complex in the cytoplasm

Although Z-DNA did not stimulate more cGAMP production in vitro, we observed increased cGAMP levels in *Tfam^+/-^* and Doxo-exposed cells, which were dependent on ZBP1 (Figures S3A and S3B). We reasoned that increased cGAS activity downstream of mtDNA stress could result from ZBP1-dependent stabilization of Z-DNA ligands for detection by cGAS and/or the redistribution of cGAS to the cytosol. Although cGAS is predominately a nuclear protein^71, 72^, we observed a significant increase in the proportion of cytosolic cGAS in ARPE-19 cells after induction of mtDNA stress by TFAM knockdown (Figures S3C and S3D). Robust cytosolic cGAS localization required ZBP1, as co-silencing of TFAM and ZBP1 led to reduced cytosolic and elevated nuclear cGAS staining, mirroring control cells (Figures S3C and S3D). Similar to TFAM-depletion, induction of mtDNA genome instability with Doxo resulted in elevated cGAS expression in cytosolic fractions (Figures S3E and S3F). To next explore whether ZBP1 could impact the distribution of cGAS independently of mtDNA stress, we transfected plasmids encoding human or mouse cGAS and ZBP1 into COS-7 cells. Consistent with the endogenous expression pattern in most cells^72^, both human and mouse cGAS localized primarily to the nucleus upon transfection (Figures S3G and 3A). In contrast, cells co-expressing cGAS and full length ZBP1 (ZBP1FL) exhibited markedly higher cytosolic intensity of cGAS, which co-localized with ZBP1 (Figures 3A, 3L, S3G and S3H). Consistent with these microscopy data, we observed a robust interaction between cGAS and ZBP1 in cytosolic immunoprecipitates (Figures S3I and 3E). Mouse fibroblast fractionation experiments revealed that ZBP1 localizes to the cytosol in *Tfam^+/-^*as well as IFN-I and Doxo treated cells (Figures S3E, 3B, and 3C). Moreover, co-immunoprecipitation (co-IP) assays in *Tfam^+/-^* and Doxo-treated cells demonstrated strong interactions between cGAS and ZBP1 in cytosolic, but not nuclear, extracts (Figure 3B and 3C). In sum, these data indicate that ZBP1 interacts strongly with cGAS in the cytoplasm.

Additional co-IP experiments to identify the domains of cGAS mediating binding to ZBP1 revealed that full-length ZBP1 could interact with both the N-terminal 147 a.a. of cGAS (cGAS N-term), as well as a construct lacking the first 147 amino acids (ΔN-term) (Figure S3J), suggesting that at least two interaction sites bridge cGAS and ZBP1. To define the domains of ZBP1 necessary for interaction with cGAS, we first generated ZBP1 constructs lacking the first (ΔZα1) or both Z-nucleic acid binding domains (ΔZα1Zα2) (Figure 3D). Upon transient expression of ZBP1 and cGAS constructs in 293FT cells, we noted that the ZBP1 Zα1Zα2 mutant was attenuated in binding cytosolic cGAS compared to full length ZBP1 (Figure 3E, lanes 1 and 3). We then developed an assay where benzonase was utilized to degrade cytosolic nucleic acids before co-IP (Figure S3K). Real time qPCR analysis of mtDNA abundance revealed that benzonase treatment was effective at degrading over 99% of mtDNA (Figure S3L). Notably, ZBP1 was less efficient at pulling down cGAS after benzonase treatment (Figure S3M), mirroring the weaker interactions observed between the ZBP1 ΔZα1Zα2 mutant and cGAS (Figure 2E). As cGAS and ZBP1 have been reported to bind a variety of endogenous and exogenous nucleic acids^73–77^, we next sought to determine if ZBP1-cGAS complexes forming in response to mtDNA instability require DNA or RNA. DNase I or RNase A treatment was sufficient to degrade mtDNA or mtRNA in *Tfam^+/-^* cells, respectively (Figure 3F). However, only DNase I treatment diminished ZBP1-cGAS binding in *Tfam^+/-^* cells displaying mtDNA instability (Figure 3G). Finally, we developed a cell-free, in vitro binding assay to examine whether ZBP1 and cGAS interact directly and explore whether the interaction is potentiated by DNA. After expressing full-length mouse cGAS and ZBP1 in *E. coli* and purification by gel filtration chromatography (Figure 3H), we performed in vitro IPs in the presence or absence of Z-or B-DNA (Figure 3I). Strikingly, these experiments revealed that ZBP1 and cGAS bind directly and that the ZBP1-cGAS complex is stabilized by DNA. Despite increased ZBP1-cGAS interactions in the presence of DNA, we observed no differences in cGAMP production after addition of recombinant ZBP1, suggesting that ZBP1 does not augment the enzymatic activity of cGAS (Figure S3N). Overall, these data constitute the novel finding that Z-DNA binding domains of ZBP1 nucleate DNA-dependent interactions with cGAS.

**Figure 3.**
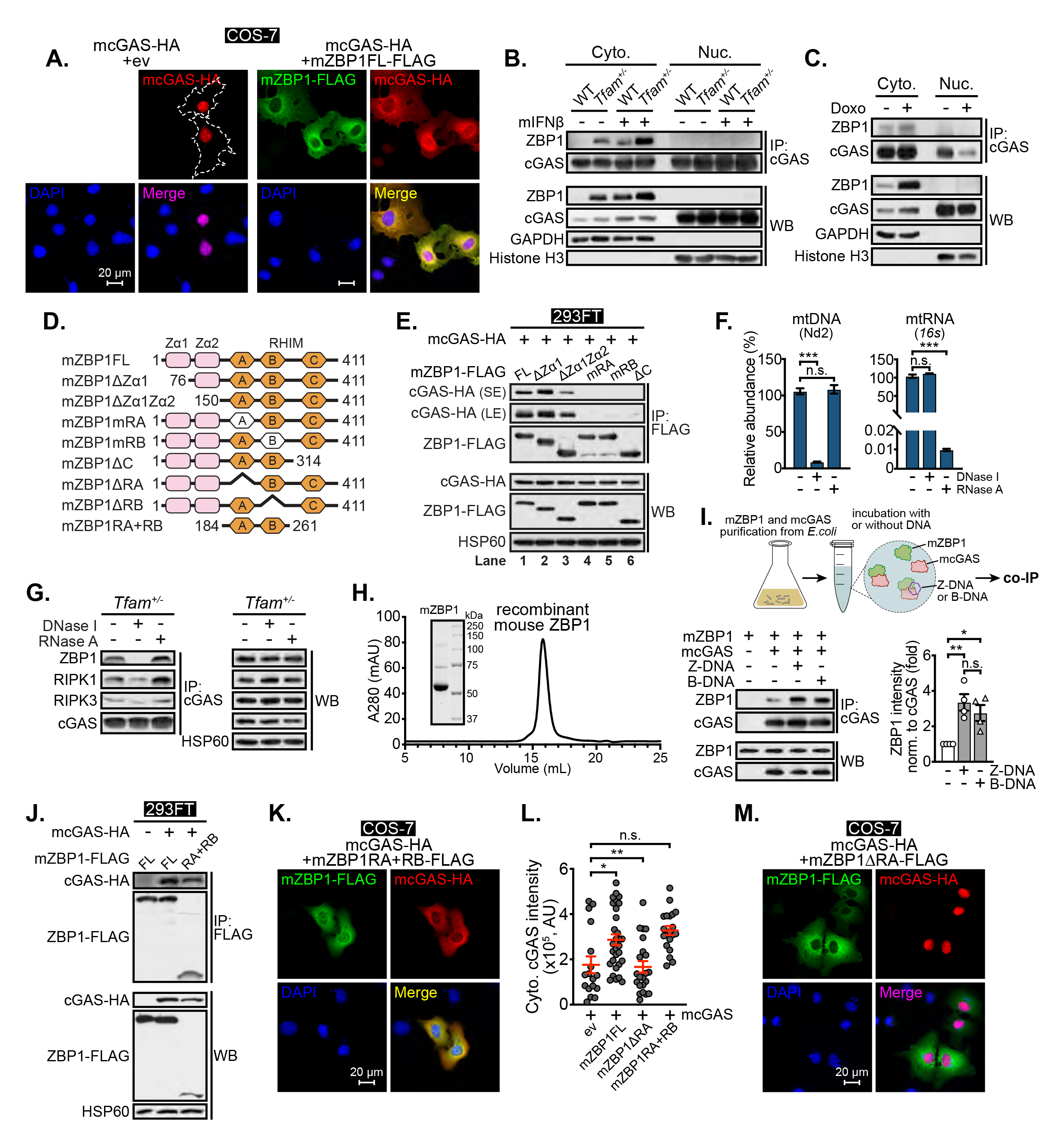
ZBP1 binds to cGAS in a DNA-and RHIM-dependent manner. **A**, Representative images of COS-7 expressing indicated proteins and stained with anti-HA and -FLAG antibodies and DAPI (ev, empty vector). **B**, Endogenous cytosolic and nuclear co-immunoprecipitations (co-IP) in WT and *Tfam^+/-^* MEFs with or without mouse IFNβ (mIFNβ) treatment (10 ng/mL, 6 h). **C**, Cytosolic and nuclear co-immunoprecipitation in WT MEFs with or without Doxo (50 nM, 48 h). **D**, Schematic representation of mouse ZBP1 full-length (FL), truncations and mutants. **E**, co-IP showing interactions between mouse cGAS and mouse ZBP1 FL, truncations or mutants in 293FT. SE, short exposure; LE, long exposure. **F**, qPCR analysis (n = 3) of mt-Nd2 abundance (left) or qRT-PCR analysis (n = 3) of mt-*16s* abundance (right) in cell lysates from **G**. **G**, Endogenous co-IP in *Tfam^+/-^* MEFs with or without DNase (100 μg/mL) or RNase (50 μg/mL) treatment. **H**, Gel-filtration chromatography and SDS–PAGE analyses of recombinant mouse ZBP1 protein. **I**, Schematic illustration of co-IP between recombinant mouse cGAS and mouse ZBP1 with or without DNA (top). co-IP showing interactions between mouse cGAS and mouse ZBP1 with or without Z-DNA or B-DNA (bottom). Densitometry of ZBP1 in cGAS immunoprecipitates is shown in graph (n = 4). Protein and DNA are mixed at molar ratio of 1:2. **J**, co-IP showing interactions between mouse cGAS and mouse ZBP1 FL and RHIMA + RHIMB fragment (RA+RB) in 293FT. **K-M**, Representative images of COS-7 expressing indicated proteins and stained with anti-HA and -FLAG antibodies and DAPI (**K, M**). Cytosolic cGAS-HA intensity is quantified in (**L**) (n ≥ 30 fields for each group. AU, arbitrary unit). Statistical significance was determined using analysis of variance (ANOVA) and Tukey post hoc test (**F, L**). *P < 0.05, **P < 0.01, and ***P < 0.001. Error bars represent SEM. See also Figure S3.

As our in vitro IP experiments revealed that ZBP1 and cGAS can bind directly in the absence of DNA, we reasoned that other domains of ZBP1 might contribute to binding. In agreement, we noted that ZBP1 constructs possessing mutated RHIM domains (mRA, mRB) (Figure 3E, lanes 4-5), lacking RHIM A or RHIM B entirely (ΔRA, ΔRB) (Figure S3O), or lacking a 97 a.a. C-terminal region containing the third RHIM domain (ΔC) (Figure 3E, lane 6) weakly interacted with cGAS upon transient expression in 293FT cells. Moreover, a ZBP1 construct of roughly 100 amino acids containing only the RHIM A and B domains (RA+RB) was able to bind cGAS (Figure 3J). Cells expressing cGAS and ZBP1RA+RB exhibited increased cytosolic cGAS and protein co-localization similar to those expressing ZBP1FL (Figures 3K and 3L). However, cGAS staining was significantly more nuclear when co-expressed with the ZBP1ΔRA mutant (Figures 3L and 3M), further supporting the notion that the cGAS and ZBP1 complex forms in the cytosol and is mediated, at least in part, by the ZBP1 RHIM domains. Taken together, our data reveal a novel cytosolic complex of ZBP1-cGAS that is stabilized by both the Z-DNA binding motifs and RHIM domains of ZBP1.

### RIPK1 and RIPK3 localize to the ZBP1-cGAS complex and sustain IFN-I signaling to mitochondrial genome instability

We next sought to clarify the mechanisms by which ZBP1 amplifies IFN-I signaling downstream of mitochondrial genome instability and Z-DNA liberation. Similar to plasmid transfection results (Figure 2I), we noted that expression of ISGs early after TFAM silencing was similar between WT and *Zbp1^-/-^*cells (Figure 4A). Notably, IFIT3 and STAT1 were comparably induced in WT and *Zbp1^-/-^* fibroblasts 72 hours post transfection with TFAM siRNAs, where the IFN-I response was first apparent (Figure 4A, lanes 4 and 10). We also observed similar levels of TBK1^S172^ and STAT1^Y701^ phosphorylation between WT and *Zbp1^-/-^* cells at the 72 hour timepoint, suggesting that initial detection of cytosolic mtDNA by cGAS, activation of STING/TBK1, and the early production of IFNβ are independent of ZBP1. However, *Zbp1^-/-^*cells displayed markedly reduced IFIT3 and STAT1 expression 96 hours post TFAM silencing (Figure 4A, lanes 6 and 12), revealing that ZBP1 is required to sustain IFN-I signaling downstream of mtDNA instability. We also observed that ZBP1 is necessary for phosphorylation of STAT1 on serine 727 at late timepoints after TFAM silencing, noting a near ablation of phospho-STAT1^S727^ in *Zbp1^-/-^*cells 96 hours after TFAM knockdown.

**Figure 4.**
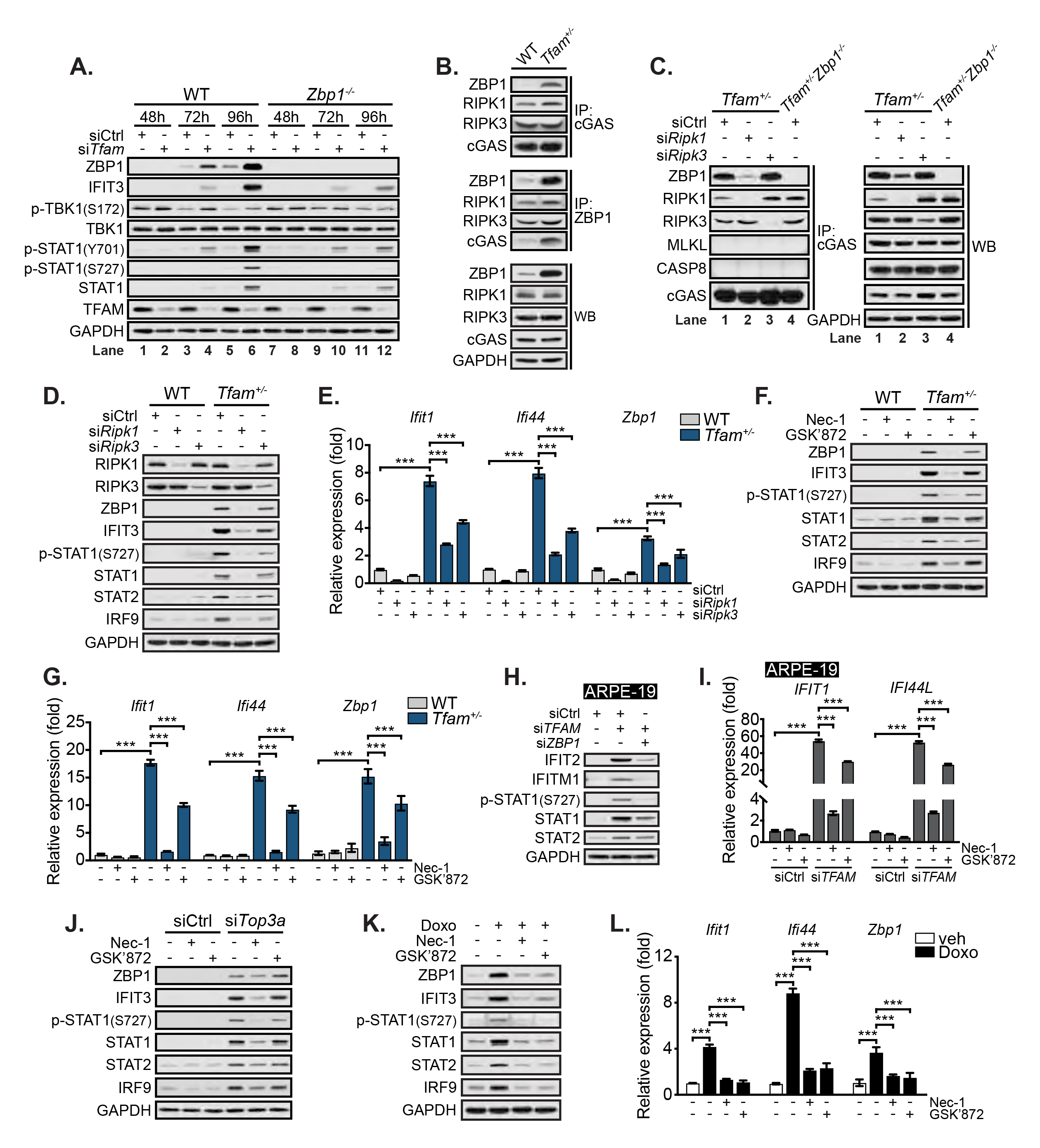
RIPK1 and RIPK3 localize to the ZBP1-cGAS complex and sustain IFN-I signaling to mitochondrial genome instability. **A**, Western blots of WT and *Zbp1^-/-^* MEFs transfected with siCtrl or si*Tfam* for indicated time. **B**, Endogenous co-IP in WT and *Tfam^+/-^* MEFs. **C**, Endogenous co-immunoprecipitations (co-IP) in *Tfam^+/-^* MEFs transfected with siCtrl, si*Ripk1* or si*Ripk3* for 48 h and *Tfam^+/-^Zbp1^-/-^* MEFs. **D, E**, Western blots (**D**) and qRT-PCR (n = 3) analysis (**E**) of WT and *Tfam^+/-^* MEFs transfected with siCtrl, si*Ripk1* or si*Ripk3* for 72 h. **F, G**, Western blots (**F**) and qRT-PCR (n = 3) analysis (**G**) of WT and *Tfam^+/-^* MEFs treated with Necrostatin-1 (Nec-1, 10 μM) or GSK’872 (2.5 μM) for 48 h. **H**, Western blots of ARPE-19 transfected with siCTRL, si*TFAM* or si*TFAM* + si*ZBP1* for 72 h. **I**, qRT-PCR (n = 3) analysis of ISGs in ARPE-19 transfected with siCTRL or si*TFAM* for 24 h and then treated with Nec-1 (100 μM) or GSK’872 (5 μM) for an additional 48 h. **J**, Western blots of WT MEFs transfected with siCtrl or si*Top3a* for 48 h and then treated with Nec-1 (10 μM) or GSK’872 (2.5 μM) for an additional 48 h. **K, L**, Western blots (**K**) and qRT-PCR (n = 3) analysis (**L**) of WT MEFs treated with vehicle or Doxo (50 nM) for 24 h and then treated with Nec-1 (10 μM) or GSK’872 (2.5 μM) for an additional 24 h. Statistical significance was determined using analysis of variance (ANOVA) and Šídák (**E, G, I, L**). *P < 0.05, **P < 0.01, and ***P < 0.001. Error bars represent SEM. See also Figure S4.

Our prior data revealed that both RIPK1 and RIPK3 localize to the DNase-sensitive ZBP1- cGAS complex in *Tfam^+/-^* cells (Figure 3G), and therefore we reasoned that RIPK recruitment/activation in the ZBP1-cGAS complex might sustain ISG expression. Interestingly, endogenous IP studies using a cGAS antibody showed that RIPK1 and RIPK3 form a complex with cGAS in WT cells and interact with ZBP1 upon its induction in *Tfam^+/-^* MEFs (Figure 4B, top panel). IPs using a ZBP1 antibody revealed that the small amount of ZBP1 in WT cells can also form a complex with RIPK1 and RIPK3 that minimally includes cGAS; however, we noted that the ZBP1- cGAS-RIPK1-RIPK3 complex was significantly enriched in *Tfam^+/-^* MEFs (Figure 4B, middle panel). Moreover, we observed that knockdown of RIPK1 in *Tfam^+/-^* cells was sufficient to reduce total ZBP1 levels and therefore the amount of ZBP1 immunoprecipitating with cGAS, whereas silencing of RIPK3 had no effect on total ZBP1 or abundance of the ZBP1-cGAS complex (Figure 4C, lanes 2 and 3). cGAS also robustly immunoprecipitated both RIPK1 and RIPK3 in *Tfam^+/-^Zbp1^-/-^* MEFs (Figure 4C, lane 4). Finally, transfection of Z-form DNA resulted in ZBP1 recruitment to the cGAS complex and augmented the level of RIPK1 co-immunoprecipitating with both proteins (Figure S4A). Collectively, these data suggest a model in which RIPK1 and RIPK3 form basal complexes with both DNA sensors, followed by the assembly of larger ZBP1-cGAS-RIPK1-RIPK3 complexes in the presence of Z-DNA ligands, ultimately culminating in the activation of RIPK signaling functions.

Phosphorylation of serine 727 in the transactivation domain of STAT1 is required to fully activate its transcriptional activity and is induced by multiple kinases including p38 MAPK, CDK8, and RIPK1^78–81^. Consistent with a role for RIPK1, we found that RIPK1 silencing dramatically reduced p-STAT1^S727^ levels and ISG expression in *Tfam^+/-^* fibroblasts, whereas RIPK3 silencing had more modest effects (Figures 4D and 4E). RIPK1 and RIPK3 inhibitor studies using Necrostatin-1 (Nec-1) and GSK’872 substantiated siRNA studies and revealed that the kinase activity of RIPK1, and RIPK3 to a lesser extent, governs p-STAT1^S727^ levels and sustain ISG expression in MEFs exhibiting mtDNA instability (Figures 4F and 4G). Additional experiments confirmed that ZBP1 and RIPK1 kinase activity are essential for STAT1^S727^ phosphorylation and ISG expression in human cells after TFAM silencing, whereas RIPK3 kinase activity is less so (Figures 4H and 4I). RIPK1 and RIPK3 kinase activities were also required to sustain p-STAT1^S727^ levels and ISG expression after silencing of TOP3A (Figures 4J and S4B) or TOP1MT (Figure S4C), as well as following exposure to Doxo (Figures 4K and 4L. Moreover, we observed that plasmid Z-DNA more potently induced IFIT3 in WT cells, likely due to increased STAT1 phosphorylation at serine 727. However, consistent with our ISG expression data (Figure 2I), both Z-and B-DNA induced comparable p-STAT1^S727^ and similar ISG expression in *Zbp1^-/-^* cells (Figure S4D). Importantly, additional experiments with Nec-1 and GSK’872 revealed that the kinase activities of RIPK1 and RIPK3 are not required for IFN-I responses triggered by canonical cGAS-STING signaling (Figure S4E), RLR-MAVS signaling (Figure S4F), or cGAS-STING signaling induced by IMM herniation and release of B-form mtDNA (Figure S4G). Taken together, these data show that following induction of mitochondrial genome stress and Z-DNA exposure, the ZBP1-cGAS complex promotes activation of RIPK1 and RIPK3, which augment STAT1^S727^ phosphorylation, increase expression and transcriptional activity of the interferon-stimulated gene factor 3 (ISGF3) complex, and sustain ISG expression.

ZBP1 is a well-appreciated driver of RIPK3 and mixed lineage kinase domain-like pseudokinase (MLKL)-dependent necroptosis, an inflammatory form of cell death that can be induced by interferon and can feed forward to potentiate IFN-I and pro-inflammatory responses^15, 82–86^. To examine the possibility that MLKL promotes mtDNA instability and release in *Tfam^+/-^* cells, we silenced MLKL by siRNA. Notably, *Tfam^+/-^* MEFs depleted of MLKL continued to show mitochondrial nucleoid aggregation and elevated ISG expression (Figure S4H-S4J). Moreover, WT and *Tfam^+/-^*MEFs were similarly susceptible to necroptosis triggered by TNFα+zVAD+CHX treatment (Figure S4K). Cell death triggered by Doxo was also equivalent between WT and *Zbp1^-/-^* MEFs (Figure S4L). In sum, these results suggest that the ZBP1-cGAS complex senses mtDNA genome instability and engages RIPK1 and RIPK3 to sustain IFN-I signaling independently of MLKL and a feed forward loop of cell death.

### Doxorubicin induces a ZBP1-dependent IFN-I response in cardiomyocytes

*Top1mt^-/-^* mice exhibit increased mtDNA instability, mitochondrial dysfunction, and cardiac injury after Doxo^87^. Consistent with our findings in fibroblasts, we also noted increased ISG expression in neonatal CMs after TOP1MT knockdown, which was entirely dependent on ZBP1 (Figure S5A). Early work in rodents reported the preferential accumulation of cardiac mtDNA damage after Doxo that lingered for weeks after administration^44^. Given the high mtDNA content in cardiomyocytes (CMs), we hypothesized that Doxo might trigger mtDNA genome instability and cytosolic release in CMs, leading to elevated ZBP1-cGAS signaling and IFN-I responses. We first mined two publicly available microarray datasets of CMs challenged with Doxo^88, 89^. Gene set enrichment analysis showed that Doxo treatment increased expression of innate immune and inflammatory genes, including those associated with IFN-I, TNFα, and DNA damage signaling (Figures 5A and S5B). Furthermore, analysis of 261 differentially expressed genes across all datasets indicated that IFN-I signaling (Interferon Alpha Response) was the most heavily enriched network (Figure 5A). In line with these bioinformatic analyses, Doxo strongly upregulated type I interferons, ZBP1, and other ISGs in human induced pluripotent stem cell-derived CMs (hiPSC-CMs) and murine neonatal CMs in a time dependent fashion (Figures 5B and 5C). Treatment of CMs with IFNAR blocking antibody further confirmed that IFN-I, but not interferon gamma or NF-κB signaling, was responsible for Doxo-induced ISG expression (Figures 5D and 5E). Suggestive of a role for Doxo in promoting mitochondrial stress, exposure of hiPSC-CMs to Doxo caused pronounced mtDNA nucleoid aggregation, mitochondrial network hyperfusion, and the accumulation of cytosolic DNA (Figure 5F). Moreover, Doxo treatment increased the intensity of Z-DNA in CMs, which was significantly reduced in cells depleted of ZBP1 (Figures 5G, 5H, and S5C). ZBP1 silencing also attenuated expression of Doxo-induced ISGs in hiPSC-CMs (Figure 5I), and Doxo failed to induce robust ISG expression in *Ifnar^-/-^, Sting^-/-^,* and *Zbp1^-/-^*neonatal CMs (Figure 5J). Collectively, these results indicate that Doxo promotes mtDNA genome instability in CMs, leading to Z-DNA accumulation that triggers the ZBP1-cGAS-IFN-I signaling axis.

**Figure 5.**
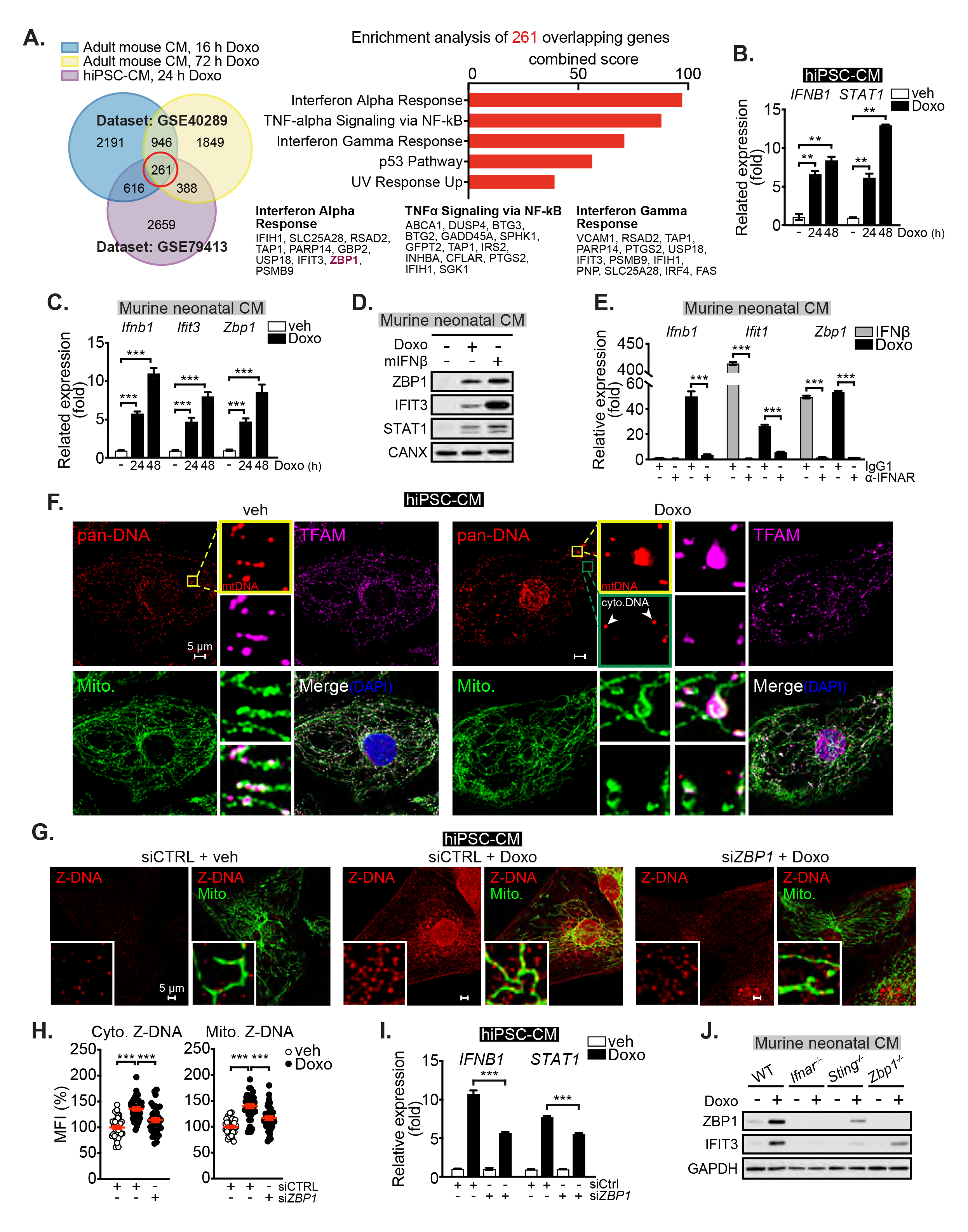
Doxorubicin induces mtDNA instability, mitochondrial Z-DNA accumulation, and IFN-I responses in cardiomyocytes. **A**, Venn diagram of differentially expressed genes overlapping across datasets GSE40289 and GSE79413 (left). The top 5 pathways identified from the 261 commonly expressed genes from all three datasets (right). Shared gene lists from the top 3 pathways are shown (bottom). **B, C**, qRT-PCR analysis (n = 3) of type I interferon transcripts and ISGs in vehicle (veh) or Doxorubicin (Doxo, 250 nM) treated human induced pluripotent stem cell-derived cardiomyocytes (hiPSC-CM) (**B**) and murine neonatal CM (**C**). **D**, Western blots of ISGs in Doxo (500 nM, 24h) or mouse IFNβ (mIFNβ, 1 ng/mL, 4 h) treated murine neonatal CMs. **E**, qRT-PCR analysis (n = 3) of *Ifnb1* and ISGs in anti-IgG1 or -interferon receptor 1 (α-IFNAR) antibody-blocked murine neonatal CMs exposed to mIFNβ or Doxo. CMs were pre-treated with anti-IgG1 or -IFNAR antibody at 15 μg/mL for 5 h before subjecting to mIFNβ (1 ng/mL, 4 h) or Doxo (500 nM, 24h) treatment. mIFNβ was used to validate the efficacy of α-IFNAR blocking antibody. **F**, Representative images of hiPSC-CMs treated with vehicle or Doxo and stained with anti-DNA, -TFAM and -PDH (Mito.) antibodies and DAPI. Inset panels are magnified 7x. **G, H**, Representative images of hiPSC-CMs transfected with siCTRL, or si*ZBP1* for 48 h, treated with vehicle or Doxo (500 nM) for an additional 24 h and stained with anti-Z-DNA and -PDH (Mito.) antibodies (**G**). Inset panels are magnified 12x. Mean fluorescent intensity (MFI) quantification of cytosolic (Cyto.) and mitochondrial (Mito.) Z-DNA is quantified in (**H**) (n ≥ 30 fields for each group from n = 3 experiments). MFI percentiles are normalized to veh-treated hiPSC-CMs. **I**, qRT-PCR analysis (n = 3) of *IFNB1* and *STAT1* in hiPSC-CMs transfected with siCTRL or si*ZBP1* for 24 h and then treated with veh or Doxo (250 nM) for an additional 48 h. **J**, Western blots of ISGs in Doxo (500 nM, 24 h) treated WT, *Ifnar^-/-^*, *Sting^-/-^* and *Zbp1^-/-^* murine neonatal CMs. Statistical significance was determined using unpaired t-test (**B, C, E**) or analysis of variance (ANOVA) and Tukey post hoc test (**H, I**). *P < 0.05, **P < 0.01, and ***P < 0.001. Error bars represent SEM. See also Figure S5.

Clinical case reports have linked IFNα immunotherapy to cardiomyopathy and other serious cardiac complications^90–95^, yet the direct effects of IFN-I on CM function have not been characterized. Interestingly, we observed that hiPSC-CMs are highly responsive to IFN-I (Figure S5D). Interestingly, IFN-I treatment increased expression of markers of CM injury (Figure S5E), while reducing expression of mtDNA-encoded OXPHOS genes (Figure S5F). Reduced mtDNA-encoded OXPHOS gene expression was also observed in murine neonatal CMs exposed to IFNβ or Doxo (Figure S5G). Using a real-time imaging assay to measure hiPSC-CM beat frequency and morphological appearance, we observed that both IFN-I and Doxo treatment shortened CM beat periodicity and induced significant vacuolization after prolonged exposure (Figure S5H). Together, these results indicate that Doxo-induced mtDNA instability triggers a ZBP1-dependent IFN-I response that potentiates CM dysfunction.

### Doxorubicin induces ZBP1 and IFN-I responses in cardiomyocytes and cardiac myeloid cells in vivo

To next determine the cellular sources and immediate kinetics of cardiac IFN-I after in vivo Doxo challenge, we employed a high dose, acute exposure protocol and assessed gene expression in CMs by RNA-seq and IFN-I activation in non-CM populations by flow cytometry (Figure 6A). Expression profiling of highly enriched CMs extracted from Doxo-exposed hearts^96, 97^ uncovered a striking upregulation of immune and IFN-I signatures mirroring our in vitro studies (Figure 5). Pathway analysis revealed ‘Interferon Alpha Response’ as the most significantly up-regulated node in CMs post-Doxo challenge (Figure 6B), with *Zbp1* representing the most highly expressed ISG (Figure 6C and Table S2). Analysis of the top downregulated pathways revealed strong inhibition of metabolic and myogenesis-related genes, with ‘Oxidative Phosphorylation’ representing the most inhibited node in CMs from Doxo-challenged hearts relative to vehicle controls (Figure 6D). Numerous genes functioning in mitochondrial oxidative metabolism were downregulated in CMs after Doxo (Figure 6E and Table S2), which is consistent with the well-appreciated mitochondrial toxicity of Doxo. Western blotting of purified CMs confirmed our RNA-seq results and showed elevated ZBP1 and IFIT3 along with reduced OXPHOS subunits ATP5A1, SDHB, and NDUFB8 from Doxo-exposed hearts relative to vehicle controls (Figure S6A). Utilizing a tamoxifen inducible, CM-specific Cre recombinase expressing strain (α*MHC-MerCreMer*) crossed onto *Sting^fl/fl^* and *Ifnar^fl/fl^* mice (Figures S6B), we observed that ablation of the cGAS-STING pathway or IFN-I signaling reduced ZBP1 expression in cardiac lysates, and other ISGs to a lesser extent (Figure S6C). This indicates that CMs in vivo are capable of directly sensing Doxo-induced DNA damage via STING and responding to IFN-I, but also reveals that other cardiac cell types contribute to IFN-I responses in the Doxo-exposed heart.

**Figure 6.**
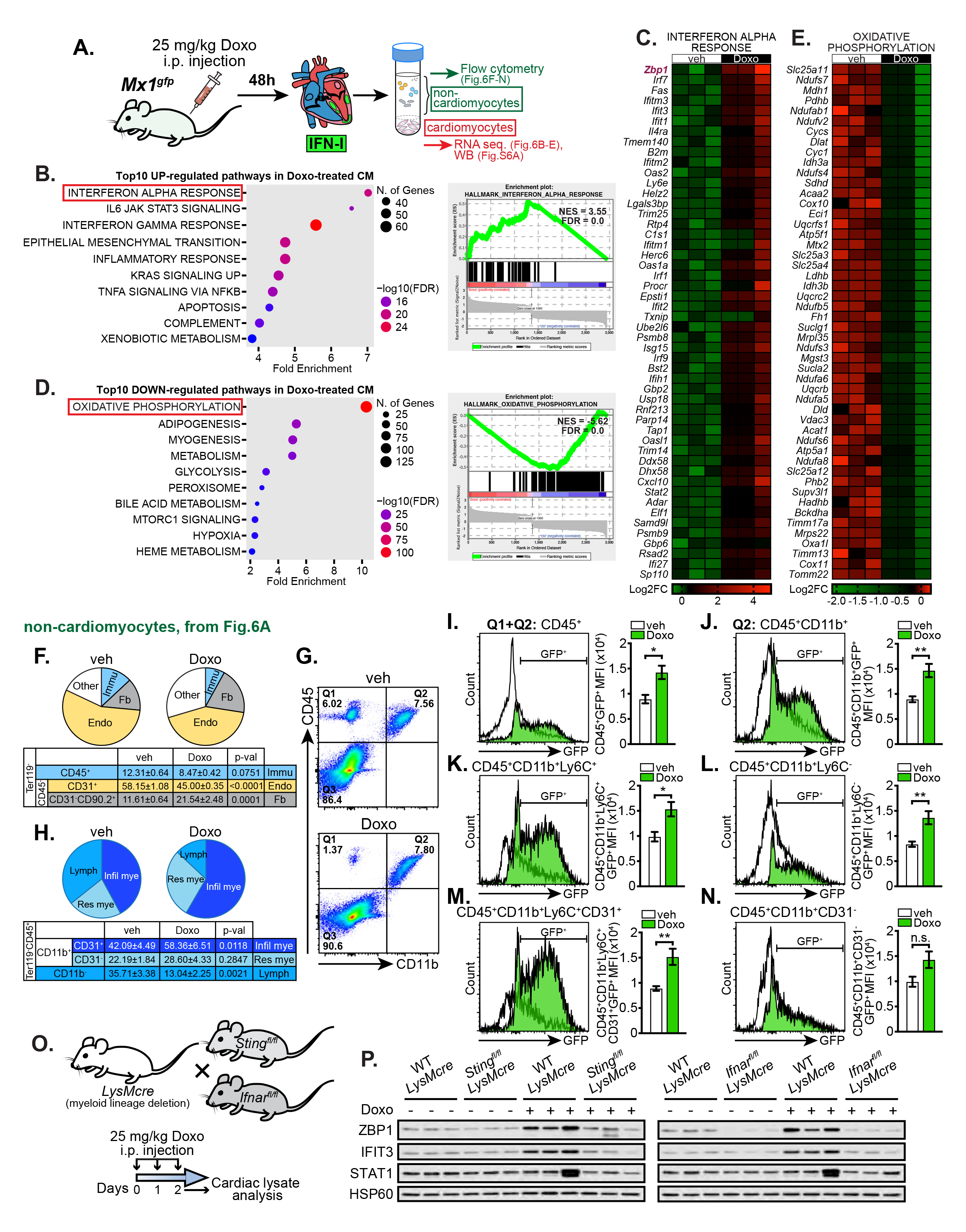
Doxorubicin induces ZBP1 and IFN-I responses in cardiomyocytes and cardiac myeloid cells in vivo. **A**, Doxorubicin (Doxo) i.p. challenge regimen, cardiac cell isolation and applications diagram. **B, D**, Pathway analysis of RNA-seq data from cardiomyocytes isolated from adult mouse hearts post Doxo challenge compared to vehicle (veh) controls. Dot plots (left) and enrichment plots (right) were generated from ShinyGo and Gene Set Enrichment Analysis (GSEA) software, respectively. NES = Normalized Enrichment Score, FDR = False Discovery Rate. **C, E**, Heatmaps of RNA-seq data displaying the top 50 up-regulated genes in interferon alpha response pathway (**C**) and top 50 down-regulated genes in oxidative phosphorylation pathway (**E**). Log2 fold changes (Log2FC) are normalized to the average of veh challenged CMs. **F**, Pie charts of major non-cardiomyocyte (non-CM) populations: immune cells (Immu), endothelial cells (Endo) and fibroblasts (Fb), compositions in vehicle (veh) and Doxo challenged mouse hearts (n = 3 per treatment). **G**, Flow cytometric analysis of CD45 and CD11b on non-CM isolated from veh and Doxo challenged mouse hearts. **H**, Pie charts of major immune cell populations: infiltrating myeloid cells (Infil mye), resident myeloid cells (Res mye) and lymphocytes (Lymph), in veh and Doxo challenge mouse hearts (n = 4 per treatment). **I-N**, Flow cytometric analysis of Mx1^gfp+^ cells in indicated cardiac immune subpopulations. Histograms are representative of 4 independent experiments. Mean fluorescent intensity (MFI) quantifications are shown on the right. **O**, Generation of myeloid-specific STING and IFNAR knockout mouse strains (top) and Doxo administration regimen (bottom). **P**, Western blots of ISGs in cardiac lysates from WT*LysMcre*, *Sting^fl/fl^LysMcre* and *Ifnar^fl/fl^LysMcre* mice post vehicle or Doxo challenge (n = 3 per genotype per challenge) (right). Statistical significance was determined using unpaired t-test (**I-N**). *P < 0.05, **P < 0.01, and ***P < 0.001. Error bars represent SEM. See also Figure S6 and Table S2.

To complement our enriched CM RNA-seq data and identify non-myocyte populations contributing to IFN-I responses in the heart after Doxo challenge, we utilized a *Mx1^gfp^* ISRE reporter mouse that expresses GFP under control of the IFN-I-regulated, endogenous *Mx1* locus^98^. We focused our antibody panels on abundant non-CM populations including Ter119^-^CD45^+^ immune cells, Ter119^-^ CD45^-^CD31^+^ endothelial cells, and Ter119^-^CD45^-^CD31^-^CD90.2^+^ fibroblasts^99^. Using these multiplex panels, we observed that Doxo promoted remodeling in all non-myocyte cardiac cell populations, with a notable decrease in endothelial cells and an increase in fibroblasts (Figure 6F). Although CD45^+^CD11b^-^ cardiac lymphoid populations decreased after Doxo, the percentage of CD45^+^CD11b^+^ myeloid immune cells remained constant (Figures 6G). However, CD45^+^CD11b^+^ cells expressing platelet/endothelial cell adhesion molecule 1 (PECAM-1; CD31), which governs leukocyte transmigration through vascular endothelial cells and recruitment into inflamed tissues, were increased in Doxo-exposed hearts (Figure 6H)^100, 101^. Despite significant changes in cardiac CD45^-^ cell numbers and CD45^+^CD11b^-^ lymphocyte cell abundance, Mx1^gfp^ intensity was not altered in either population after Doxo treatment (Figures S6D and S6E). However, Mx1^gfp^ expression was markedly increased in CD45^+^CD11b^+^ cardiac myeloid cells (Figures 6I and 6J) comprised of Ly6C^+^ monocytes (Figure 6K) and Ly6C^-^ macrophages and dendritic cells (Figure 6L)^102^. Most Ly6C^+^ monocytes exhibiting high Mx1^gfp^ also expressed CD31 in Doxo-exposed hearts relative to vehicle controls (Figure 6M). In contrast, CD11b^+^CD31^-^ cells did not exhibit significant increases in Mx1^gfp^ (Figure 6N), indicating that recruited myeloid cells are largely responsible for non-CM IFN-I signaling post-Doxo. Consistent with a role for myeloid immune cells in the cardiac IFN-I program induced by Doxo, conditional ablation of STING (*Sting^fl/fl^LysMcre*) or IFNAR (*Ifnar^fl/fl^LysMc*re) in myeloid cells (Figure 6O) markedly reduced ZBP1 and other ISGs in cardiac lysates (Figure 6P). Finally, we observed that ZBP1 is upregulated in cardiac extracts and interacts strongly with cGAS after in vivo Doxo challenge (Figure S6F). Collectively, these data suggest that Doxo induces significant cardiac remodeling, robust ZBP1-cGAS complex assembly, and induction of IFN-I responses in vivo.

### Adjuvant IFN**α** synergizes with Doxorubicin to enhance cardiac and mitochondrial dysfunction in tumor-bearing mice

In order to expand our in vitro findings and advance mechanistic understanding of IFN-I-mediated cardiotoxicity, we merged an autochthonous melanoma model^103, 104^ with an extended Doxo protocol that more closely resembles clinical timelines in late DIC (Figure S6G). Doxo chemotherapy significantly restricted melanoma growth, with adjuvant IFNα providing a modest synergistic effect (Figure S6H). Transthoracic echocardiographic analyses four weeks after cessation of Doxo chemotherapy revealed reduced left ventricular ejection fraction (EF), fractional shortening (FS) and left ventricle (LV) mass in tumor-bearing mice (Figure S6I). This was accompanied by significant left ventricle dilation, wall thinning, and cardiac fibrosis (Figures S6J-S6L). Consistent with our acute Doxo challenge model, we noted that ZBP1 and other ISGs were elevated in Doxo-exposed hearts relative to mice receiving vehicle only (Figure S6M). Strikingly, mice receiving Doxo plus adjuvant IFNα exhibited more pronounced left ventricular dysfunction, elevated cardiac fibrosis, and higher cardiac ISG expression compared to the Doxo only group (Figures S6I-S6M). Finally, melanoma bearing mice receiving Doxo and IFNα showed reduced OXPHOS protein expression and cardiac mtDNA abundance (Figures S6N and S6O). These results suggest that IFN-I signaling synergizes with Doxo to promote loss of mtDNA and OXPHOS subunit expression, which correlate with increased cardiac fibrosis and heart failure.

### ZBP1 is a key regulator of cardiac IFN-I responses that contribute to Doxorubicin-induced cardiomyopathy

As our data indicate that multiple cardiac populations contribute to IFN-I responses that potentiate Doxo-induced cardiac dysfunction, we next employed a chronic Doxo dosing regimen on ZBP1, STING, and IFNAR constitutive knockouts (Figure 7A). Echocardiography of WT mice revealed marked impairments in cardiac functional measures of left ventricular EF and FS after Doxo (Figure 7B). In contrast, mice lacking ZBP1, STING or IFNAR exhibited higher EF and FS compared to WT throughout the longitudinal study (Figure 7B). In depth analysis of longitudinal echocardiographic measurements showed that *Zbp1^-/-^*, *Sting^-/-^* and *Ifnar^-/-^* mice displayed less Doxo-induced wall thinning, left ventricle dilation, and weight loss compared to WT controls (Figures 7C, 7D and S7A-S7C). To link cardiac dysfunction with sustained IFN-I signaling, we examined transcript and protein abundance of ISGs seven weeks after the cessation of Doxo injection. WT mice exhibited sustained expression of ISGs assayed at week 10 post-Doxo. In contrast, ZBP1 and other ISGs were significantly lower or not induced in Doxo-exposed *Zbp1^-/-^*, *Sting^-/-^* and *Ifnar^-/-^* mice (Figures 7E and S7D). Moreover, all knockout strains exhibited reduced cardiac expression of gene signatures associated with heart failure (Figure 7F). In agreement with cardiac transcript expression, histologic analyses of Doxo-challenged *Zbp1^-/-^*, *Sting^-/-^* and *Ifnar^-/-^* mice revealed significantly less cytoplasmic vacuolization at week 10 compared to WT controls (Figures 7G and S7E), and all knockout strains had markedly lower cardiac fibrosis as revealed by picrosirius red staining (Figures 7H and S7F). Finally, Doxo-induced OXPHOS subunit loss as measured by immunofluorescence staining for Complex V subunit ATP5A1 was significantly lessened in *Zbp1^-/-^*, *Sting^-/-^* and *Ifnar^-/-^* mice compared to WT animals (Figures 7I and S7G). Additional analysis of *Sting^-/-^*and *Ifnar^-/-^* cardiac lysates revealed higher levels of mtDNA and OXPHOS subunit expression after Doxo relative to WT controls (Figures S7H and S7I). In summary, our data reveal that the ZBP1-cGAS-STING axis governs cardiotoxic IFN-I signaling, which contributes significantly to the pathobiology of chemotherapy-related heart failure.

**Figure 7.**
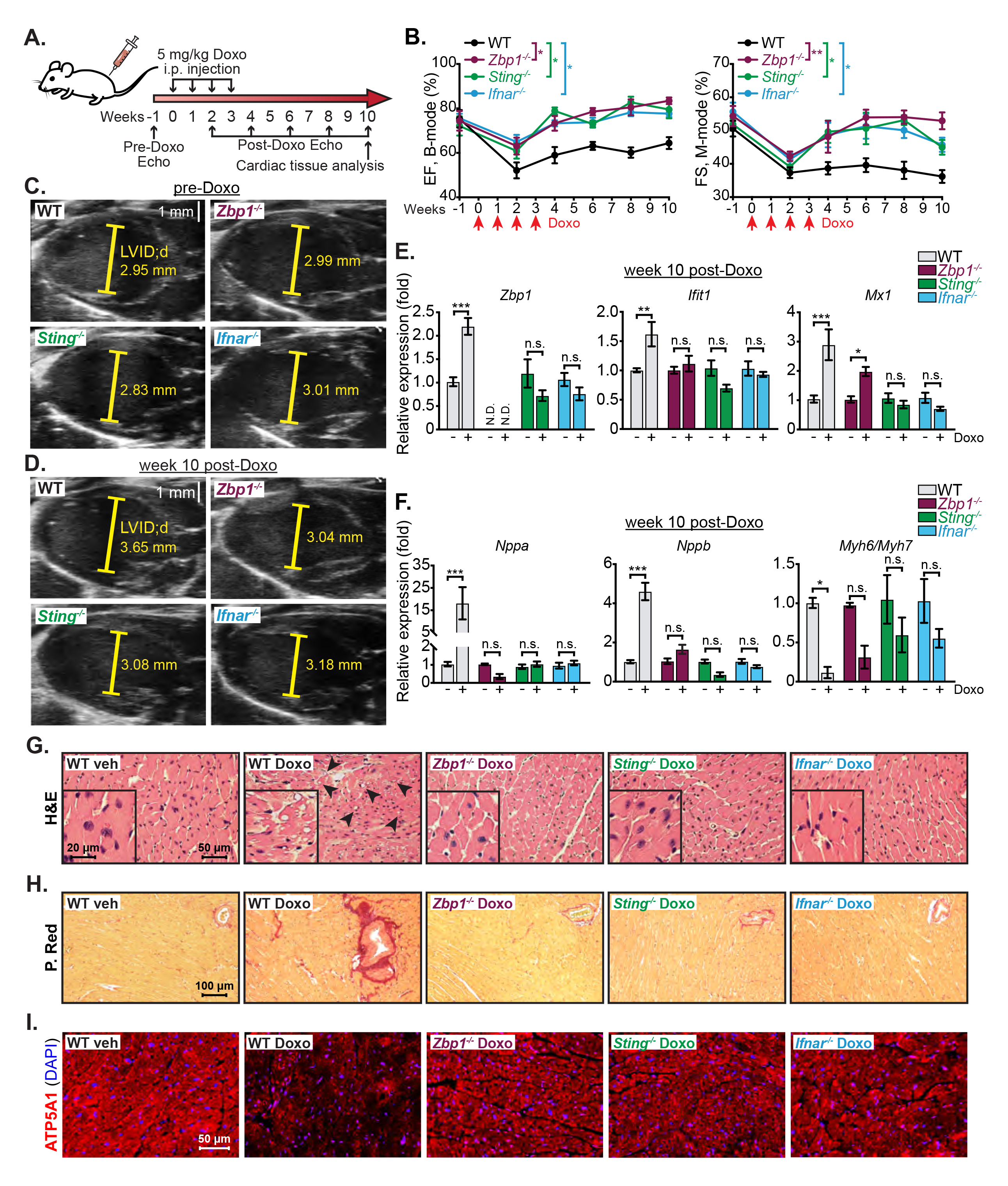
The ZBP1-IFN-I axis contributes to Doxorubicin-induced cardiac injury and cardiomyopathy. **A**, Doxorubicin (Doxo) i.p. challenge regimen, longitudinal cardiac monitoring, and cardiac harvest timepoint diagram. **B**, Left ventricular ejection fraction (EF) and fractional shortening (FS) calculated from B-mode or M-mode images using Vevo Lab software (n = 6 per genotype). **C, D**, Representative parasternal long axis images from each genotype pre-Doxo (**C**) and at week 10 post Doxo challenge (**D**). **E, F**, qRT-PCR analysis of ISGs (**E**) or natriuretic peptides *Nppa*, *Nppb* and the *Myh6/7* expression ratio (**F**) in vehicle (veh) or Doxo treated WT, *Zbp1^-/-^*, *Sting^-/-^* and *Ifnar^-/-^* cardiac lysates (n = 3 per genotype per treatment). **G-I**, Representative images of H&E staining (**G**), picrosirius red (P. Red) staining (**H**) and ATP5A1 and DAPI immunofluorescent staining (**I**) of the heart sections from veh treated WT mice and 10-week post Doxo treated WT, *Zbp1^-/-^*, *Sting^-/-^* and *Ifnar^-/-^* mice. Arrows in **G** indicate significant cardiomyocyte vacuolization in Doxo-treated WT heart section. Statistical significance was determined using analysis of variance (ANOVA) and Dunnett (**B**) or Šídák (**E, F**) post hoc test. *P < 0.05, **P < 0.01, and ***P < 0.001. Error bars represent SEM. See also Figure S7.

## DISCUSSION

ZBP1/DLM-1/DAI was initially identified as a cytosolic DNA sensor that activates NF-κB and IFN-I signaling^105, 106^. However, experiments in *Zbp1^-/-^* cells and mice revealed that ZBP1 is dispensable for IFN-I and pro-inflammatory cytokine induction to exogenous and endogenous B-form DNA^14, 16, 49^. Subsequent work showed that ZBP1 predominately senses endogenous and microbial RNA, leading to inflammatory responses and necroptotic cell death^14, 15, 17–20^. More recent genetic and biochemical studies uncovered a key role for ZBP1 in the autoinflammatory pathology of mice lacking the RNA editing enzyme Adenosine deaminase acting on RNA (ADAR1), which accumulate endogenous retroelement-derived, dsRNA in the left-handed Z conformation^16, 21, 22, 73^. ZBP1 can also sense nuclear Z-form DNA^73^, and therefore ZBP1 has emerged as a key PRR for double-stranded nucleic acids in the Z-conformation. Our study highlights that ZBP1 also plays a central role in orchestrating innate immune responses to mitochondrial genome instability and the subsequent release of topologically abnormal, Z-form mtDNA. Although a computational study identified Z-prone sequences in the non-coding region of mtDNA^107^, our study is the first to report that genetic and pharmacologic triggers of mtDNA stress induce Z-DNA in cellulo. As a circular genome, mtDNA is susceptible to over accumulation of positive and negative supercoils that form ahead of and behind mitochondrial replication and transcription complexes^108^. Increased negative mtDNA supercoiling in cells lacking TFAM, TOP3A, and TOP1MT favors B-Z transition and mitochondrial stress, leading to the release of mitochondrial Z-DNA. Since overall Z-DNA intensity was significantly lower in *Zbp1^-/-^* cells, we posit that the Zα domains of ZBP1 stabilize Z-DNA to sustain detection by cGAS and augment IFN-I signaling. Our data reinforce this hypothesis, as we found that transfection of plasmid DNA in the Z-confirmation elicited significantly higher ISGs than plasmid B-DNA in WT, but not *Zbp1^-/-^*, cells. Overall, our data provide strong evidence that mitochondrial Z-DNA is immunostimulatory and suggest that ZBP1 stabilizes mitochondrial Z-DNA and/or brings it into a complex with cGAS to sustain IFN-I responses.

Biochemical studies revealed that the Zα and RHIM domains of ZBP1 are required for maximal cytosolic interactions with cGAS. ZBP1 forms homotypic interactions with the RHIM domain-containing proteins RIPK1 and RIPK3 to mediate NF-κB activation and innate immune signaling, as well as promote RIPK3-MLKL-dependent necroptosis^15, 109, 110^. However, MLKL silencing did not impact IFN-I responses in cells experiencing mtDNA stress. Therefore, we conclude that the ZBP1-cGAS complex functions to sustain the IFN-I program downstream of mitochondrial Z-DNA release, independently of necroptotic cell death. ZBP1, RIPK1, and RIPK3 can induce a subset of ISGs in mouse cortical neurons during ZIKA virus infection^111^, and human ZBP1 induces cell death-independent inflammatory signaling via RIPK1 and RIPK3^112^. Consistent with immune signaling roles for RIPK1 and RIPK3, we find that both kinases are present in the ZBP1-cGAS complex and are required to sustain IFN-I signaling and ISG expression downstream of mitochondrial Z-DNA liberation. Thus, our data are in line with a recent study showing that ZBP1 interacts with cGAS, RIPK1, and RIPK3 in *Casp8^-/-^/Mlkl^-/-^* MEFs harboring Z-RNA^113^. Notably, the kinase activities of RIPK1, and RIPK3 to a lesser extent, promote STAT1^S727^ phosphorylation in response to mitochondrial Z-DNA accumulation and are required to sustain the ZBP1-dependent IFN-I program. Future studies should determine whether CASP8, FADD, and/or MLKL activities are inhibited in cells experiencing mitochondrial genome instability, which might favor ZBP1-cGAS-RIPK signaling to STAT1 while repressing ZBP1-dependent cell death signaling.

We also observed that mtDNA stress strongly upregulates ZBP1 and feedforward IFN-I signaling in cardiac cells and tissue. Importantly, we have uncovered that Doxo induces mtDNA aggregation and Z-DNA accumulation in CMs, which trigger a robust IFN-I signature that is dependent on ZBP1, STING, and IFNAR. Doxo and other anthracycline chemotherapeutics are among the most effective and widely used antineoplastic drugs, however their clinical application is limited by adverse cardiac side effects that occur in many patients^114^. We find that CMs are exquisitely sensitive to IFN-I, which promotes expression of cardiac damage markers, induces functional changes in beat periodicity, and reduces expression of mtDNA-encoded OXPHOS transcripts. Our data also reveal that both CMs and infiltrating CD31^+^CD11b^+^ myeloid cells respond to Doxo by activating ZBP1- dependent IFN-I responses. Using a Doxo dosing protocol that closely resembles clinical timelines in late DIC, we find that ZBP1 and other ISGs remain elevated in the heart for seven weeks after the last Doxo dose, suggestive of a self-propagating IFN-I cycle. Consequently, ZBP1, STING, and IFNAR knockout mice are significantly protected from DIC and display marked improvements in LV function and cardiac output. We therefore conclude that the ZBP1-cGAS axis is a key innate immune pathway that propagates cardiotoxic IFN-I responses and enhances cardiac remodeling and heart failure after Doxo exposure.

In conclusion, we have uncovered that ZBP1 is a key sensor of mitochondrial Z-DNA that works cooperatively with cGAS to sustain IFN-I responses in vitro and in vivo. Mechanistically, ZBP1 complexes with cGAS, RIPK1, and RIPK3 in the cytosol and amplifies cGAS activity in response to mtDNA stress via its ability to sense and stabilize cytosolic Z-form DNA. Although cGAS and IFN-I signaling have recently been implicated in heart failure^28, 31, 115–117^, a role for ZBP1 has not been explored. Our findings indicate the presence of a ZBP1-dependent IFN-I signaling program in cardiac cells and tissues exposed to Doxo and highlight that ZBP1 is a key innate immune regulator of cardiotoxicity and heart dysfunction. ZBP1 may therefore represent a novel therapeutic target in chemotherapy-related cardiac dysfunction, as well as other cardiovascular disorders where mtDNA stress sustains disease-promoting IFN-I and inflammatory responses.

## LIMITATIONS OF THE STUDY

Although we observe increased Z-DNA staining in mitochondria and the cytosol after induction of mitochondrial genome instability, we did not identify the precise Z-DNA sequences that are stabilized by ZBP1. Mitochondrial dysfunction and mtDNA replication defects can feed forward to cause nuclear genome instability^118, 119^, and therefore it is possible that Z-DNA derived from both mtDNA and nuclear DNA accumulate in our models. Second, although we find that mitochondrial genome instability upregulates ZBP1 and promotes ZBP1-cGAS interactions, both sensors form basal complexes with RIPK1 and RIPK3. Our study does not elucidate the stoichiometry of these complexes, nor identify the molecular changes that specifically license RIPK activity for increased STAT phosphorylation and IFN-I responses, but not cell death. Finally, we show that mitochondrial DNA instability predominantly triggers ZBP1-cGAS-RIPK-dependent IFN-I signaling in mouse and human cells, yet we cannot exclude additional roles for ZBP1-driven cell death in DIC. Future studies will hopefully clarify these limitations and promote the development of ZBP1-focused therapies for Doxo-related heart injury.

## ACKNOWLEDGEMENTS

We thank Dr. Ken Ishii and Dr. Bill Kaiser for providing *Zbp1^-/-^* mice; R. Moore for flow cytometry assistance; and members of the Shadel, Upton, Li, and West labs for helpful discussions and feedback on the manuscript. This work was supported by NIH grants to A.P.W. (R01HL148153 and R01CA193522), C.W.T. (R01HL145534), P.L. (R01AI145287), G.S.S. (R01AR069876 and R01CA216101) and J.W.U. (R21AI135709). Additional support was provided by NIH grant P30ES029067, and a Natural Sciences and Engineering Research Council of Canada grant to T.E.S. (NSERC - RGPIN-2016-04083). G.S.S. is the Audrey Geisel Chair in Biomedical Science. Y.L. was supported by an American Heart Association Predoctoral Fellowship (Grant 825908). S.T.-O. was supported by a Ruth L. Kirschstein National Research Service Award (NRSA) Predoctoral Fellowship to Promote Diversity in Health-Related Research (F31HL160141).

## AUTHOR CONTRIBUTIONS

Y.L. and A.P.W. designed the experiments, analyzed the data, and wrote the manuscript. Y.L. carried out most of the in vivo and in vitro assays, performed statistical analyses, and generated figures. J.J.V. assisted with immunoprecipitation and immunofluorescence assays. Y.C. assisted with transcriptional analysis of hiPSC-CMs, cardiac cell isolation, flow cytometry, and animal experiments. J.D.B. assisted with B-DNA and Z-DNA generation and in vivo Doxo challenge. Y.Li., B.W. and P.L assisted with cGAS and ZBP1 protein purification, cGAS activity assays, and provided expertise on cGAS studies. S.T.-O. assisted with cell culture assays. D.F. and M.S. assisted with immunofluorescence labeling and image collection from cardiac sections. L.C.W. performed melanoma induction and tumor monitoring. K.B.R. and J.W.U. provided ZBP1 plasmids and advised on ZBP1 assays. J.D., A.M., and T.E.S. assisted with mtDNA southwestern blots and provided expertise on TOP1MT. O.N.B. and S.D.Y. assisted with immunoprecipitation assays. C.W.T. provided expertise and advice on cardiac measurements. M.W.B. provided the inducible melanoma model and advice on tumor induction and monitoring. G.S.S. provided the *Tfam^+/-^*mouse model and guidance on mtDNA assays. A.P.W. conceived the project, performed RNA-seq processing and bioinformatic analyses, and provided overall supervision. All authors read and approved the manuscript.

## INCLUSION AND DIVERSITY

We worked to ensure sex balance in the selection of non-human subjects. One or more of the authors of this paper self-identifies as an underrepresented ethnic minority in their field of research or within their geographical location. One or more of the authors of this paper self-identifies as a gender minority in their field of research. One or more of the authors of this paper self-identifies as living with a disability. One or more of the authors of this paper received support from a program designed to increase minority representation in their field of research.

**Figure S1. ZBP1 is required for robust IFN-I responses downstream of mtDNA stress, related to Figure 1. A**, qPCR analysis (n = 3) of mtDNA abundance in the cytosol (Cyto.) and mitochondrial (Mito.) fraction from WT and *Tfam^+/-^* MEFs. **B,** Heatmaps of RNA-seq data displaying up-regulated DNA and RNA sensors in WT and *Tfam^+/-^* MEFs. Fold changes (FC) are normalized to the average of WT MEFs. **C,** qRT-PCR analysis (n = 3) of ISGs in WT, *Sting^-/-^* and *Mavs^-/-^* MEFs transfected with si*Tfam* for 72 h. **D,** qRT-PCR analysis (n = 3) of ISGs in WT and *Tfam^+/-^* MEFs transfected with siCtrl or si*Zbp1* for 96 h. **E,** Southwestern blots of mtDNA isolated from WT and *Tfam^+/-^* MEFs (left). Quantification of supercoiled to relaxed mtDNA ratio (n = 3, right). **F,** qRT-PCR analysis (n = 3) of *Top3a* in WT, *cGAS^-/-^* and *Zbp1^-/-^* MEFs transfected with siCtrl or si*Top3a* for 96 h. **G,** qRT-PCR analysis (n = 3) of *TOP3A* in ARPE-19 transfected with siCTRL or si*TOP3A* for 72 h. **H,** qPCR analysis (n = 3) of mtDNA abundance in WT MEFs transfected with siCtrl or si*Top3a* with or without 2’,3’-dideoxycytidine (ddC, 100 μM) for 72 h. **I, J,** qRT-PCR analysis (n = 3) of *Top1mt* (**I**) and ISGs (**J**) in WT, *cGAS^-/-^* and *Zbp1^-/-^* MEFs transfected with siCtrl or si*Top1mt* for 96 h. **K,** qPCR analysis (n = 3) of mtDNA abundance in WT MEFs transfected with siCtrl or si*Top1mt* with or without ddC (100 μM) for 72 h. **L,** qRT-PCR analysis (n = 3) of ISGs in WT MEFs transfected with siCtrl or si*Top1mt* with or without ddC (100 μM) for 72 h. **M,** qPCR analysis (n = 3) of mtDNA abundance in Cyto. and Mito. fraction from vehicle (veh) or Doxorubicin (Doxo) treated WT MEFs. **N,** qRT-PCR analysis (n = 3) of ISGs in Doxo (125 nM) treated MEFs. **O,** Western blots of ISGs in Doxo (48 h) treated MEFs. **P,** qRT-PCR analysis (n = 3) of ISGs in WT, *Ifnar^-/-^* and *Mavs^-/-^* MEFs treated with or without Doxo (500 nM, 24 h). **Q, R,** qPCR analysis (n = 3) of mtDNA abundance (**Q**) and qRT-PCR analysis (n = 3) of ISGs (**R**) in WT MEFs treated with veh or Doxo (250 nM, 24 h) with or without ddC (100 μM) for 72 h. **S,** qRT-PCR analysis (n = 3) of ISGs in Doxo (50 nM, 48 h) treated WT and *Zbp1^-/-^* MEFs with or without IFNβ priming (10 pg/mL, 18 h). **T, U,** qRT-PCR analysis (n = 3) of ISGs in WT and *Zbp1^-/-^* MEFs transfected with ISD (2 μg/mL) for 6 h (**T**) or treated with ABT-737 (ABT, 10 μM) + Q-VD-OPH (QVD, 10 μM) for 6 and 24 h (**U**). Statistical significance was determined using unpaired t-test (**A, C, E, F, G, I, M, P**), or analysis of variance (ANOVA) and Tukey (**D, H, J, K, L, N, Q, R**) or Šídák (**S, T, U**) post hoc test. *P < 0.05, **P < 0.01, and ***P < 0.001. Error bars represent SEM.

**Figure S2. Mitochondrial genome instability promotes the accumulation and cytosolic release of mitochondrial Z-form DNA that potentiates IFN-I responses, related to Figure 2. A**, Quantification of unshifted DNA band from 3 independent electrophoretic mobility shift assay (EMSA) experiments in Fig. 2B. **B,** Slot blots of B-DNA and Z-DNA. **C,** Representative images of WT MEFs transfected with polyethylenimine (PEI) complexed B-DNA or Z-DNA and stained with anti-Z-DNA antibody and SYTOX. **D,** Schematic illustration of DNA extraction and transfection from different subcellular pools (left). Slot blots of cytosolic, mitochondrial, and nuclear DNA isolated from WT MEFs treated with vehicle (veh) or Doxorubicin (Doxo). **E,** Representative images of WT MEFs transfected with si*Top3a* or si*Top1mt* or treated with Doxo with or without ddC (100 μM). Anti-pan-DNA and -HSP60 (Mito.) antibodies were used for staining. Inset panels are magnified 4x. **F,** Mean fluorescent intensity (MFI) quantification of cytosolic (Cyto.) and mitochondrial (Mito.) Z-DNA fluorescent intensities in WT MEFs transfected with si*Top3a* or si*Top1mt* or treated with Doxo with or without ddC (100 μM) (n ≥ 30 fields for each group from n=3 experiments). MFI percentiles are normalized to groups without ddc treatment. **G,** Quantification of Cyto. and Mito. Z-DNA MFI in WT MEFs treated with ABT-737 (ABT, 1 μM) + Q-VD-OPH (QVD, 1 μM) for 6 and 24 h. MFI percentiles are normalized to DMSO control-treated (0 h) WT MEFs. **H,** Quantification of Cyto. and Mito. Z-DNA fluorescent intensities in WT MEFs transfected with siCtrl or si*Tfam* for 72 h and treated with Hydralazine (Hyd, 100 μM) for 24 h (n ≥ 20 fields for each group from n = 3 experiments). MFI percentiles are normalized to WT MEFs transfected with siCtrl. **I,** qRT-PCR analysis (n = 3) of ISGs in WT and *Zbp1^-/-^* MEFs transfected with siCtrl or si*Tfam* for 72 h and treated with or without Hydralazine (100 μM) for 24 h. **J,** qRT-PCR analysis (n = 3) of ISGs in ARPE-19 transfected with B-DNA or Z-DNA (1 μg/mL) for 16 h with or without IFN-I priming (10 pg/mL, 8 h). **K,** Enzyme activity assay of cGAS in the presence of B-DNA, Z-DNA, salmon sperm DNA or no DNA. Statistical significance was determined using analysis of variance (ANOVA) and Tukey (**A, H**), Dunnett (**G**) or Šídák (**I, J**) post hoc test, or unpaired t-test (**F**). *P < 0.05, **P < 0.01, and ***P < 0.001. Error bars represent SEM.

**Figure S3. ZBP1 interacts with cGAS in the cytoplasm, related to Figure 3. A**, cGAMP production in WT and *Tfam^+/-^* MEFs (n ≥ 4) measured by ELISA. **B,** cGAMP production in Doxo-treated WT and *Zbp1^-/-^* MEFs (n = 8) measured by ELISA. **C, D,** Representative images of ARPE-19 transfected with siCTRL, si*TFAM* or si*TFAM* + si*ZBP1* for 72 h and stained with anti-cGAS and -DNA antibodies and DAPI (**C**). Inset panels are magnified 4x. Cytosolic (Cyto.) and nuclear (Nuc.) fractions of cGAS are quantified in (**D**) (n ≥ 30 fields for each group. AU, arbitrary unit) **E, F,** Western blots of cytosolic and nuclear proteins in vehicle (veh) or 50 nM Doxorubicin (Doxo) treated WT MEFs (**E**). cGAS intensities from post-nuclear extract (PNE) and nuclear extract (NE) in veh or Doxo treated WT MEFs are quantified in (**F**). 2 different MEF lines are shown in the western blots, and 3 different MEF lines were used for quantification. **G, H,** Representative images of COS-7 expressing indicated proteins and stained with anti-HA and -EGFP antibodies and DAPI (**G**). Cytosolic cGAS-HA intensity is quantified in (**H**) (n ≥ 30 fields for each group). **I,** co-IP showing interactions between human ZBP1 and human cGAS in 293FT. **J,** Schematic representation of the mouse cGAS full-length (FL) and truncations (top). co-IP showing interactions between mouse ZBP1 and mouse cGAS FL and truncations in 293FT (bottom). **K,** Schematic illustration of co-IP method used in **M**. **L,** qPCR analysis (n = 3) of mt-ND1 abundance in cell lysates from **M**. **M**, co-IP between mouse cGAS and mouse ZBP1 in 293FT with or without benzonase treatment. **N,** Enzyme activity assay of cGAS in the presence of B-DNA or Z-DNA, with or without ZBP1. **O,** co-IP showing interactions between mouse cGAS and mouse ZBP1 FL and deletions in 293FT. Statistical significance was determined using unpaired t-test (**A, B, F, H, L**), or analysis of variance (ANOVA) and Tukey post hoc test (**D**). *P < 0.05, **P < 0.01, and ***P < 0.001. Error bars represent SEM.

**Figure S4. RIPK1 and RIPK3 mediate mtDNA stress-induced IFN-I signaling independently of MLKL and cell death, related to Figure 4. A**, co-IP from WT MEFs transfected with Z-DNA or B-DNA (100 ng/mL, 24 h). **B, C,** qRT-PCR analysis (n = 3) of ISGs in WT MEFs transfected with siCtrl, siTop3a (**B**) or siTop1mt (**C**) for 48 h and then treated with Nec-1 (10 μM) or GSK’872 (2.5 μM) for an additional 48 h. **D,** Western blots of WT and *Zbp1^-/-^* MEFs transfected with B-DNA or Z-DNA (125 ng/mL) for 24 h. **E-G,** qRT-PCR analysis (n = 3) of ISGs in WT MEFs transfected with ISG (2 μg/mL) for 12 h (**E**), Poly I:C (1 μg/mL) for 6 h (**F**), or treated with ABT-737 (ABT, 10 μM) + Q-VD-OPH (QVD, 10 μM) for 6 h (**G**). **H,** Representative fluorescence images of WT and *Tfam^+/-^* MEFs transfected with siCtrl and si*Mlkl* for 96 h and stained with anti-DNA and -HSP60 antibodies and DAPI. Inset panels are magnified 4x. **I, J,** qRT-PCR analysis (n = 3) (**G**) and western blots (**H**) of MLKL and ISGs in WT and *Tfam^+/-^*MEFs transfected with siCtrl or si*Mlkl* for 96 h. **K,** LDH assay (n = 4) showing cytotoxicity in WT and *Tfam^+/-^* MEFs after 8 h of TNFα (25 ng/mL) + z-VAD-FMK (zVAD) (50 μM) + cycloheximide (CHX) (5 μg/mL) treatment. **L,** LDH assay (n = 3) showing cytotoxicity in WT and *Zbp1^-/-^* MEFs after 48 h of 500 nM Doxo treatment. Statistical significance was determined using analysis of variance (ANOVA) and Tukey (**I**) or Šídák (**B, C, E, F, G**) post hoc test, or unpaired t-test (**K, L**). *P < 0.05, **P < 0.01, and ***P < 0.001. Error bars represent SEM.

**Figure S5. IFN-I and Doxorubicin synergize to imbalance cardiomyocyte mitochondrial gene expression and disrupt contractility, related to Figure 5. A**, qRT-PCR analysis (n = 3) of *Top1mt* and ISGs in WT and *Zbp1^-/-^* murine neonatal cardiomyocytes (CMs) transfected with siCtrl or si*Top1mt* for 96 h. **B,** Volcano plots of the Gene Set Enrichment Analysis (GSEA) showing the differentially regulated pathways in GSE40289 and GSE79413 datasets: cardiomyocytes (CM) isolated from adult mouse hearts challenged with 25 mg/kg Doxorubicin (Doxo) for 16 h vs saline controls, and human induced pluripotent stem cell-derived cardiomyocytes (hiPSC-CM) treated with 1 μM Doxo for 24 h vs controls (NES = Normalized Enrichment Score, FDR = False Discovery Rate). **C,** Mean fluorescent intensity (MFI) quantification of cytosolic (Cyto.) and mitochondrial (Mito.) Z-DNA in WT and *Zbp1^-/-^* murine neonatal CMs treated with or without Doxo (125 nM, 24 h) (n ≥ 30 fields for each group). MFI percentiles are normalized to untreated WT or *Zbp1^-/-^* murine neonatal CMs. **D-F,** qRT-PCR analysis (n = 3) of ISGs (**D**), cardiac damage markers (**E**) and mitochondrial DNA-or nuclear DNA-encoded mitochondrial transcripts (**F**) in hiPSC-CMs treated with or without human recombinant IFNα (10 ng/mL) and IFNβ (10 ng/mL) for 96 h. **G,** qRT-PCR analysis (n = 3) of mitochondrial DNA-encoded mitochondrial transcripts in murine neonatal CMs treated with mouse IFNβ (1 ng/mL) or Doxo (500 nM) for 24 h. **H,** Camera-capture assay showing beat periodicity and images of hiPSC-CMs after indicated treatments (100 nM Doxo and/or 10 ng/mL of both human recombinant IFNα and IFNβ) for 96 h and 120 h. Yellow arrows indicate cardiomyocyte vacuolization. AU, arbitrary unit. Statistical significance was determined using analysis of variance (ANOVA) and Tukey post hoc test (**A, C**), or unpaired t-test (**D-G**). *P < 0.05, **P < 0.01, and ***P < 0.001. Error bars represent SEM.

**Figure S6. Doxorubicin induces cardiomyocyte IFN-I signaling that potentiates cardiomyopathy in tumor bearing mice, related to Figure 6. A**, Western blots of ISGs and OXPHOS components in cardiomyocytes (CM) isolated from vehicle (veh) or Doxorubicin (Doxo) challenged hearts (n = 7 per challenge). **B,** Generation of tamoxifen (TAM)-inducible cardiomyocyte-specific STING and IFNAR knockout mouse strains (left) and Doxo administration regimen (right). **C,** Western blots of ISGs in cardiac lysates from WT*MCM*, *Sting^fl/fl^MCM* and *Ifnar^fl/fl^MCM* mice induced with TAM and then challenged with vehicle or Doxo (n = 3 per genotype per challenge) (bottom). **D, E,** Flow cytometric analysis of GFP^+^ population in CD45^-^ (**D**) or CD45^+^CD11b^-^ (**E**) cells isolated from veh and Doxo challenged mouse hearts. Histograms are representative of 4 independent experiments. Mean fluorescent intensity (MFI) quantifications are shown on the right. **F,** co-IP showing interactions between ZBP1 and cGAS in cardiac extracts from veh and Doxo-challenged mice. Mice were i.p. injected with 25 mg/kg Doxo for 24 hrs. **G,** Schematic illustration of melanoma induction. **H,** Primary melanoma volume of each treated group measured every other week starting from 5 weeks post 4- Hydroxytamoxifen (4OHT) induction. Mice were i.p. injected with 5 mg/kg Doxo once a week for 4 weeks. Recombinant mouse IFNα was administered twice weekly for 4 weeks via i.p. injection of 20,000 U per dose. **I,** Left ventricular ejection fraction (EF), fractional shortening (FS) and mass at week 4 were calculated from parasternal long axis B-mode. (n = 5 per group). **J, K,** Representative parasternal long axis (**J**) and short axis (**K**) images from each treatment at week 6 post vehicle, Doxo or Doxo+IFNα challenge. **L,** Representative picrosirius red staining images of heart sections from vehicle-, Doxo-and Doxo+IFNα treat melanoma mice. Collagen area of each group are quantified and shown at up-right corner. **M,** Western blots (n = 2) of ISGs in cardiac extracts from vehicle, Doxo and Doxo+IFNα challenged tumor-bearing mice at week 8. **N, O,** Western blots of mitochondrial OXPHOS components (n = 2) (**N**) and qPCR analysis (n = 3) of mtDNA abundance (**O**) in cardiac extracts from veh, Doxo and Doxo+IFNα challenged tumor-bearing mice at week 8. Statistical significance was determined using unpaired t-test (**D, E, O**), or analysis of variance (ANOVA) and Tukey post hoc test (**H, I, L**). *P < 0.05, **P < 0.01, and ***P < 0.001. Error bars represent SEM.

**Figure S7. Doxorubicin-induced ISG expression and cardiac dysfunction are lessened in the absence of ZBP1-STING-IFNAR signaling, related to Figure 7. A**, Left ventricle wall thickness and internal diameter measurements were made pre-Doxo (week −1) and at week 4 and 8 weeks post challenge in short axis M-mode (n = 12 of WT and *Zbp1^-/-^* groups, n = 6 of *Ifnar^-/-^* and *Sting^-/-^* groups). **B,** Heart weight changes in WT, *Zbp1^-/-^*, *Sting^-/-^* and *Ifnar^-/-^* mice at week 8 compared to pre-Doxo treatment (week −1) (n = 12 of WT and *Zbp1^-/-^* groups, n = 6 of *Ifnar^-/-^* and *Sting^-/-^* groups). **C,** Representative short axis images from each genotype pre-and 10 weeks post-Doxo treatment. **D,** Western blots (n = 2 per genotype per treatment) of ISGs in cardiac lysates 10 weeks post Doxo challenge. **E,** Percentages of cardiomyocytes exhibiting vacuolization in WT, *Zbp1^-/-^*, *Sting^-/-^* and *Ifnar^-/-^* heart sections with or without Doxo challenge (n = 12 fields per group). **F,** Percentages of picrosirius red positive area (collagen area) in WT, *Zbp1^-/-^*, *Sting^-/-^* and *Ifnar^-/-^* heart sections from mice with or without Doxo challenge (n = 9 fields per group). **G,** Quantification of ATP5A1 fluorescent intensity in WT, *Zbp1^-/-^*, *Sting^-/-^* and *Ifnar^-/-^* heart sections with or without Doxo challenge (n = 10 fields per group, AU, arbitrary unit). **H,** qPCR analysis (n = 3 per genotype per treatment) of mtDNA abundance in cardiac lysates from vehicle-and Doxo-treated WT, *Ifnar^-/-^*and *Sting^-/-^* mice. **I,** Western blots (n = 2 per genotype per treatment) of mitochondrial OXPHOS subunits in cardiac lysates from vehicle-and Doxo-treated WT, *Ifnar^-/-^* and *Sting^-/-^* mice. Statistical significance was determined using analysis of variance (ANOVA) and Tukey post hoc test (**A, E, F, G, H**), or unpaired t-test (**B**). *P < 0.05, **P < 0.01, and ***P < 0.001. Error bars represent SEM.

## STAR METHODS

## RESOURCE AVAILABILITY

### Lead contact

Further information and requests for resources and reagents should be directed to and will be fulfilled by the lead contact, A. Phillip West.

### Materials availability

All primary cells, plasmids, and mouse strains generated in this study are available from the Lead Contact with a completed Materials Transfer Agreement.

### Data and code availability

All data needed to evaluate the conclusions drawn herein are present in the paper and/or the supplementary figures and tables. Any additional information required to reanalyze the data reported in this paper is available from the lead contact upon request. RNA-seq datasets have been deposited in the Gene Expression Omnibus under accession numbers GSE223553 and GSE223698.

## EXPERIMENTAL MODEL AND SUBJECT DETAILS

### Mouse husbandry and strains

C57BL/6J (strain 000664), *LysMCre* (strain 004781), *αMHC-MerCreMer* (strain 005657), *Sting^fl/fl^* (strain 031670), *Ifnar1^fl/fl^* (strain 028256), *Mx1^gfp^* (strain 033219), *Sting^-/-^* (*Sting1^gt^*; strain 017537) and *Ifnar1^-/-^*(MMRRC Strain #032045-JAX) mice were purchased from The Jackson Laboratory. *Mavs^-/-^*(strain 008634) mice were purchased from The Jackson Laboratory and were backcrossed onto the C57BL/6J background for 10 generations before use. *Tfam^+/-^* mice were previously reported^46, 48^. *Zbp1^-/-^* mice^49^ were obtained collaboratively from Dr. Ken Ishii and were backcrossed onto the C57BL/6J background for at least 4 generations before use. Breeding colonies of all strains were established and maintained on the C57BL/6J background at Texas A&M University. Mice were group-housed in humidity-controlled environments maintained at 22°C on 12-hour light–dark cycles (600–1800). Food and water were available ad libitum. All animal procedures were approved by the Texas A&M Animal Care and Use Committee (TAMU IACUC). The euthanasia methods employed in the study are consistent with the recommendations of the American Veterinary Medical Association (AVMA).

### Doxorubicin administration

Male and female mice between 8-10 weeks of age were housed in a standard 12-h light/dark cycle with unlimited access to food and water. Doxorubicin stock solution (10 mg/mL) was prepared in cell culture grade water (VWR, 82007-330) and used within 28 days. Further dilutions were made in DPBS (Sigma-Aldrich, D8537) on the day of injection. In Figure 6, *Mx1^gfp^*, WT*LysMcre*, *Sting^fl/fl^LysMcre* and *Ifnar^fl/fl^LysMcre* mice were intraperitoneally (i.p.) injected with one dose of 25 mg/kg of Doxorubicin. In Figure S6, Cre expression was induced by 3 consecutive daily i.p. injections of 30 μg/g of tamoxifen into WT*αMHC-MerCreMer*, *Sting^fl/fl^αMHC-MerCreMer* and *Ifnar^fl/fl^αMHC-MerCreMer* mice, followed by a 4-week resting period, and subsequent application of one dose of 25 mg/kg of Doxorubicin via i.p. challenge. In Figure 7, intraperitoneal (i.p.) injection of Doxorubicin (5 mg/kg body weight, 100 μL injection volume) was performed weekly for four continuous weeks (cumulative dose 20 mg/kg body weight).

### Mouse melanoma induction, Doxorubicin and adjuvant IFN**α** therapy

An autochthonous model of melanoma was utilized in Figures S6H-S6P, and melanomas were induced on 21-day old mice as described in Dankort et al. 2009^103^. First, mice were anesthetized by isoflurane, then a commercial topical depilatory cream was applied to remove hair from a 2 centimeter squared region on the lower dorsal region of mice approximately 2 centimeters from the base of the tail. The next day, mice were anesthetized with isoflurane and 1 μL of a 20 mM solution of 4-hydroxytamoxifen (Hello Bio, HB6040) dissolved in ethanol was applied topically to the exposed skin. Four weeks after melanoma induction, hair was removed from the lower dorsal region of the mice using hair trimmers to allow confirmation of tumor induction and monitoring of tumor growth. Tumors were measured for width, length, and height every 2-4 days using digital calipers in order to calculate tumor volume. Mice received vehicle (PBS) or Doxorubicin (5 mg/kg body weight, 100 μL i.p. injection volume) weekly for four consecutive weeks (cumulative dose 20 mg/kg body weight). Adjuvant IFNα (20,000 units) was co-administrated together with Doxorubicin in a total volume of 100 μL or diluted with PBS in a total volume of 50 μL and i.p. injected alone as indicated in Figure S6I.

### Cardiac cell isolation

Cardiac cell isolation was performed as described by Ackers-Johnson et al.^96^ with some modifications. Briefly, *Mx1^gfp^* mice i.p. injected with saline or Doxorubicin were anaesthetized using an isoflurane vaporizer, and anesthesia was maintained using a nose cone throughout the surgical procedure. Following anesthesia, thoracic cavities were opened to expose the heart. The descending aorta was severed to allow exsanguination, and 7 mL EDTA buffer was immediately injected into the right ventricle. Next, the ascending aorta was clamped using a hemostat, and the heart was removed then transferred to a 60 mm dish containing fresh EDTA buffer, then again injected with 10 mL EDTA buffer into the left ventricle at 2 to 3 mm above the heart apex. Next, the heart was transferred to a 60 mm dish containing perfusion buffer and injected with 3 mL of perfusion buffer through the same perforation on the left ventricle. Then, the heart was transferred to a 60 mm dish containing collagenase buffer, and digestion was achieved by continuously injecting collagenase buffer through the same perforation at the flow rate approximately 2 mL/min until the heart turned pale and flaccid and the injection perforation was enlarged. The atria and ventricles were gently separated, and then the ventricles (with septum) were transferred to a new 60 mm dish containing 3 mL fresh collagenase buffer. The digested tissues were pulled into 1 mm pieces using fine-tip forceps, single cardiac cells were dissociated by gently trituration, and finally the enzyme activity was neutralized by adding 5 mL stop buffer. Cell suspension was filtered through a 70 µm strainer, and cardiomyocytes were settled by gravity for 20 min in a 50 mL tube. The supernatant containing non-myocyte cardiac cells was subjected to staining for flow cytometry analysis, and the cardiomyocyte pellet was preserved for RNA and protein extraction for further analysis. All buffers were prepared in cell-culture grade water and recipes can be found in Ackers-Johnson et al^96^, with the exception that 2,3-butanedione monoxime (BDM) was replaced with 5mM blebbistatin (Selleckchem, S7099). EDTA, perfusion, and collagenase buffers were apportioned into 10 mL syringes with 27 G needles and 60 mm dishes, then kept warm at 37 °C.

### Cell culture

Primary WT, *Tfam^+/−^*, *Zbp1^−/−^*, and *Tfam^+/-^Zbp1^−/−^* MEFs were generated from E13.5–14.5 embryos, maintained in DMEM (Sigma-Aldrich, D5796) supplemented with 10% low endotoxin FBS (VWR, 97068-085), and sub-cultured no more than five passages before experiments. Murine neonatal cardiomyocytes were isolated from 2-3 day old neonates with Neonatal Cardiomyocyte Isolation Kit (Miltenyi Biotec, 130-098-373). Human iPSC-derived cardiomyocytes were purchased from Fujifilm Cellular Dynamics (01434) and cultured as described in the manual. 293FT cells were purchased from Invitrogen (R70007). COS-7 (CRL-1651) and ARPE-19 (CRL-2302) cells were purchased from ATCC.

## METHOD DETAILS

### siRNA and DNA transfection

siRNA transfection was performed with 25 nM siRNA duplexes and 3 μL of Lipofectamine RNAiMAX reagent (ThermoFisher Scientific, 13778150) according to the manufacturer’s protocol. DNA transfection was performed with polyethylenimine, PEI (Alfa Aesar, 43896) at a ratio of 5:1 PEI to nucleic acids.

### Southwestern Blot

mtDNA isolation and southwestern blotting were performed as described previously^54^. In brief, MEFs were treated with 10 µM aphidicolin (Hello Bio, HB3690) for 4 h, after which 50 µM BrDU (Invitrogen, B23151) was added to the culture medium for an additional 24 h. Following treatment, DNA was isolated using the E.Z.N.A.® plasmid DNA minikit (Omega bio-tek, D6942-02), and 1 µg of isolated DNA was loaded and run on a 0.45% agarose gel in TBE (26 V at 4 °C for 18-20 h). After electrophoresis, the agarose gel was washed with water and soaked in denaturing buffer (0.5M NaOH, 1.5M NaCl) for an hour with gentle shaking. Afterwards, the gel was rinsed again with water and soaked in 10X SSC for 30-60 min before capillary transfer to an ImmobilonP membrane (Millipore Sigma), as described previously^120^. Once transferred, the membrane was soaked in 6X SSC before undergoing UV cross-linking (UV Stratalinker 1800). The cross-linked membrane was blocked with 10% milk in TBST for 1 h, followed by immunoblotting with anti-BrdU antibody (BD Biosciences, 555627) for 2 h at room temperature. The membrane was then washed and incubated with HRP-linked anti-mouse IgG (EMD Millipore Etobicoke) for 1 h, after which the blot was developed using Supersignal West Femto enhanced chemiluminescence substrate (ThermoFisher Scientific) and imaged with a chemiluminescence imaging analyzer (Fujifilm, LAS3000mini).

### Cloning and retroviral transduction

Murine STING-HA (InvivoGen) was subcloned into the pMXs-IRES-Puro vector and replication incompetent retroviruses were packaged using Platinum-A cells (Cell Biolabs, RV-102) according to the manufacturer’s instructions. 293FT were exposed to supernatants containing pMXs-IRES-Blast-STING-His retroviruses and 4 μg/mL polybrene (Millipore, TR-1003-G). Two days post transduction, 10 μg/mL blasticidin was added to select a stable expression population.

### Topologically constrained Z-DNA synthesis

Circular Z-DNA and B-DNA were synthesized as described by Zhang et al.^61^. Briefly, 74-mer oligos containing a 5’ phosphate modification (Sigma-Aldrich) were ligated to form circular single-stranded DNA (ssDNA) using a stepwise reaction. The 150 µL reaction mixture contained 25 µM splint, 75 U T4 ligase (ThermoFisher Scientific, EL0011), and 0.1X T4 ligase buffer. A second mix contained the corresponding 74-mer DNA oligo at 22 µM in 150 µL 0.1X T4 ligase buffer. 15 µL of the DNA mix was added to the reaction mix every 20 min for a total of 10 additions bringing the total reaction volume to 300 µL. After the final addition of DNA, the reaction was allowed to continue for another 12 h. The reaction was carried out at 20 °C and terminated by incubation at 65 °C for 10 min. Any remaining linear DNA was degraded by addition of 35 µL 10X Exonuclease I buffer and 20 µL Exonuclease I (ThermoFisher Scientific, 720735KU) and incubation at 37 °C for 2 h followed by 80 °C for 20 min. Circular ssDNA strands were purified by phenol/ chloroform extraction and ethanol precipitation.

ssDNA strands were annealed to form double-stranded B-or Z-DNA as follows. The annealing reaction contained 10 mM HEPES and 10 mM MgCl_2_ at pH 7.5. Topologically constrained Z-DNA was obtained by annealing circular 74F ssDNA to circular 74R ssDNA strands at a 1:1 molar ratio. DNA strands were annealed in a thermocycler using the following steps: 90 °C for 5 min, cool to 60 °C at 0.1 °C/s, hold at 60 °C for 10 min, cool to 20 °C at 0.1 °C/s, and hold at 12 °C. B-DNA was made by annealing linear 74F ssDNA to circular 74R ssDNA or annealing circular 74F ssDNA to linear 74R ssDNA as described above. After annealing, fully double-stranded B-DNA was obtained by incubating the dsDNA with T4 ligase in 1X T4 ligase buffer for 12 h at 25 °C followed by incubation with Exonuclease I for 2 h at 37 °C and exonuclease inactivation at 80 °C for 20 min. Both B-DNA and Z-DNA were then purified by phenol/ chloroform extraction and ethanol precipitation. Purified DNA pellets were redissolved in 10 mM HEPES buffer (10 mM MgCl_2_, pH 7.5).

### Electrophoretic Mobility Shift Assay (EMSA)

Circular Z-and B-DNA were diluted to 10 ng/µL (0.43 µM) in 10 mM HEPES buffer (10 mM MgCl_2_, pH 7.5) and incubated with anti-Z-DNA antibody (Absolute antibody, Ab00783-23.0) at a 1:1 or 1:3 DNA:antibody molar ratio for 4 h at 25 °C. DNA-antibody complexes were resolved by electrophoresis on native 10 % TBE-polyacrylamide gels (ThermoFisher Scientific, EC6275BOX). DNA was visualized by post-staining with SYBR Green I (MedChemExpress, HY-K1004) and imaging on a Bio-Rad ChemiDoc MP Imaging System.

### Protein expression and purification

Full length mouse ZBP1 and mouse cGAS were cloned into a modified pET28a vector with a N-terminal Avi-His6-Sumo tag. Both ZBP1 and cGAS were expressed in *E. coli* BL21(DE3) with 0.4 mM isopropyl β-D-1-thiogalactopyranoside (IPTG) induction overnight at 15 °C. Proteins were purified using Ni-NTA agarose (QIAGEN), followed by size-exclusion chromatography using HiLoad 16/600 Superdex200 columns (GE Healthcare Life Sciences). The Avi-His6-Sumo tag was cleaved by sumo protease and removed using Superdex200 columns. Mouse ZBP1 was eluted in the buffer containing 20 mM Tris-HCl pH 7.5, 150 mM NaCl. Mouse cGAS was eluted in the buffer containing 20 mM Tris-HCl pH 7.5, 500 mM NaCl.

### cGAS activity assay

The cGAS activity assay was performed in 20 mM HEPES pH 7.5, 150 mM NaCl, 5 mM MgCl_2_, 5 mM DTT, 2.5 mM ATP, 2.5 mM GTP. 2 µM mouse cGAS was incubated with 0.1 µg/µL B-DNA or Z-DNA with or without 2 µM mouse ZBP1 for 1 h at 37 °C. 2 µM mouse cGAS was incubated with 0.1 µg/µL salmon sperm DNA (Invitrogen, 15632-011) or no DNA for 1 h at 37 °C as the positive and negative control. Reactions were quenched by adding 20 mM EDTA. The products were filtered by 10 kDa-cutoff Amicon Ultra centrifugal filters to remove DNA and proteins and then analyzed by a Mono Q 5/50 GL column (GE Healthcare Life Sciences).

### Co-immunoprecipitation

For Figures 3B and 3C, MEFs were treated with 10 ng/mL of mouse IFNβ for 6 hrs or in 50 nM of Doxo for 48 hrs, respectively, before co-IP. The cytosolic fraction was obtained by lysing the cells with NP-40 lysis buffer (50 mM Tris pH7.5, 0.15 M NaCl, 1 mM EDTA, 1% NP-40 and 10% Glycerol), and nuclear fraction was acquired using Nuclear Complex co-IP kit (Active Motif, 54001), which allows a gentle release of DNA-bound protein complexes in the nucleus. Rabbit TrueBlot® Set (Rockland, 88-1688-31) was used for both pre-cleaning and co-IP from both fractions. Mouse TrueBlot® Set (Rockland, 88-7788-31) was used for pre-cleaning and co-IP with ZBP1 antibody in Figure 4B. For co-IPs of cGAS and ZBP1 constructs in 293FT (Figures 3E, 3J, S3I, S3J, S3M and S3O), Pierce™ Control Agarose Resin (ThermoFisher Scientific, 26150) and Anti-DYKDDDDK Tag (L5) Affinity Gel Antibody (Biolegend, 651502) were used for pre-cleaning and IP, respectively. Due to different subcellular localizations of cGAS mutants, the cytosolic fraction and nuclear fraction were obtained as described above, combined and then used for co-IPs of cGAS mutants and ZBP1 in 293FT (Figures S3J). To degrade cytosolic nucleic acids in Figure S3M before co-IP, NP-40 cell lysates were incubated with 10 U/μL of benzonase (Millipore, 71205-3) for 4 h at 4 °C. BSA was added to the samples that were not treated with benzonase to even protein concentration. 20 μL of sample was separated and the benzonase was inactivated by adding 0.5 μL of 0.5 M EDTA and heating to 95°C for 15 min. 10 ng/μL DNA was used in qPCR to amplify mtDNA using primers against mt-ND1. To degrade cytosolic DNA or RNA in Figure 3G before co-IP, NP-40 cell lysates were incubated with 100 μg/mL of DNase I (Worthington Biochemical, LS002139) or 50 μg/mL of RNase A (Worthington Biochemical, LS005650) for 4 h at 4 °C. After incubation, DNA was extracted using Phenol/chloroform and 10 ng/μL DNA was used in qPCR to amplify mtDNA using primers against mt-ND2; RNA was extracted using Quick-RNA microprep kit (Zymo Research, R1051) and subjected to qRT-PCR to amplify *16s* RNA.

### Subcellular fractionation, DNA isolation and slot blotting

3 x 10^6^ MEFs were seeded in 15 cm dishes overnight and treated with vehicle or 500 nM of Doxo for 24 hrs. Subcellular fractionation and DNA isolation were performed as previously described^121^. In brief, cells were lysed in digitonin lysis buffer (150 mM NaCl, 50 mM HEPES pH 7.4, and 5 μg/mL digitonin), incubated end over end for 10 min at 4 °C, and centrifuged at 950 x g for 5 min at 4 °C. The cytosolic supernatants were transferred into a new tube and saved as cytosolic fraction, and the pellets were lysed in NP-40 lysis buffer (50 mM Tris pH 7.5, 0.15 M NaCl, 1 mM EDTA, 1% NP-40 and 10% Glycerol). After centrifugation of NP-40 lysates at 21130 x g for 10 min at 4 °C, the supernatants were transferred and saved as mitochondrial fraction, and the nuclei pellets were further lysed in and SDS lysis buffer (20 mM Tris pH 8 and 1% SDS). SDS lysates were boiled at 95°C for 5 min and centrifuged at 21130 x g for 5 min at RT. The supernatants were transferred and saved as nuclear fraction. All fractions were treated with 50 μg/mL of RNase A for 1.5 hrs at 37°C and 200 μg/mL of Proteinase K for 1 h at 55°C before being subjected to Phenol-Chloroform based DNA isolation. After DNA isolation, 200 μg of cytosolic DNA, 1 mg of mitochondrial DNA and 50 μg of nuclear DNA were blotted onto 0.45 μm Nylon membranes (Whatman, 10416230). After air drying, membranes were incubated in primary antibodies at 4°C overnight in 1X PBS containing 1% casein, HRP-conjugated secondary antibody at room temperature for 1 h, and then developed with Luminata Crescendo Western HRP Substrate (Millipore, WBLUR0500).

### Quantitative PCR

To measure relative gene expression by qRT–PCR in cells and tissues, total cellular RNA was isolated using Quick-RNA microprep kit (Zymo Research, R1051). Approximately 0.5-1 μg RNA was isolated, and cDNA was generated using the qScript cDNA Synthesis Kit (Quanta, 95047-100). cDNA was then subjected to qPCR using PerfeCTa SYBR Green FastMix (Quanta, 84069). Technical triplicates were run each biological sample, and average expression values were normalized against *Rpl37* cDNA using the 2^-ΔΔCT^ method^121^. For mtDNA abundance assessment, 10 ng/μL of template DNA was used for qPCR, and expression values of each replicate were normalized against nuclear-encoded *Tert or RNA18S*^121^.

### Immunoblotting

Cells and tissues were lysed in NP-40 lysis buffer or SDS lysis buffer supplemented with protease inhibitor (Roche, 04693159001) and then centrifuged at 4 °C to obtain cellular lysate. After BCA protein assay (ThermoFisher Scientific, 23235), equal amounts of protein (10–40 μg) were loaded into 10-20% SDS–PAGE gradient gels and transferred onto 0.22 μM PVDF membranes (Bio-Rad, 1620177). After air drying to return to a hydrophobic state, membranes were incubated in primary antibodies at 4 °C overnight in 1X PBS containing 1% casein, HRP-conjugated secondary antibody at room temperature for 1 h, and then developed with Luminata Crescendo Western HRP Substrate (Millipore, WBLUR0500). For cGAS quantifications in Figures S3E and S3F, cytosolic cGAS expression was normalized to GAPDH, and nuclear cGAS was normalized to LMNB1. These normalized values were used to calculate the percentage of cGAS in the cytosolic (cytosolic cGAS/ total cGAS) and nuclear (nuclear cGAS/ total cGAS) compartments.

### Immunofluorescence microscopy

Cells were grown on coverslips and treated as described. After washing in PBS, cells were fixed with 4% paraformaldehyde for 20 min, permeabilized with 0.1% Triton X-100 in PBS for 5 min, blocked with PBS containing 5% FBS for 30 min, stained with primary antibodies for 1 h, and stained with secondary antibodies for 1 h. Cells were washed with PBS containing 5% FBS between each step. Coverslips were exposed to 200 ng/mL 4′,6-diamidino-2-phenylindole (DAPI) (ThermoFisher Scientific, 62247) for 2 min to label nuclei, then mounted with ProLong Diamond Antifade Mountant (Invitrogen, P36961) and sealed with clear nail polish. For experiments requiring cytosolic signal quantification, HSC CellMask Green (Invitrogen, H32714) was added to coverslips at 2 μg/mL for 30 min, then washed several times in PBS, before DAPI staining. For Figure S2C, SYTOX (Invitrogen, MP33025) was added to coverslips at 0.17 μM for 15 min, then washed in PBS. For Figure 1J, images were acquired on a LSM 780 confocal microscope (Zeiss) with a 40x oil-immersed objective and ZEN 3.3 software. For Figures 1F, 2C, 2E, 2G, S2C, S2E, 3A, 3K, 3M, S3C, S3G, S4H, 5F and 5G, images were acquired on an ECLIPSE Ti2 (Nikon) with a 60x oil-immersed objective and NIS-Elements AR 5.21.02 software. Representative images shown in Figures 1F, 2C, 2E, 2G, S2E, S3C, S4H, 5F and 5G were imported into Huygens Essential software (v21.4.0) and deconvolved using the ‘aggressive’ profile in the Deconvolution Express application. In Figures 3A, 3K, 3M, S3C and S3G, the nuclear cGAS intensity in each cell was quantified within the DAPI positive area, and cytosolic cGAS intensity per cell was quantified in the CellMask positive area minus the DAPI positive area. In Figures 2D, 2F, 2H, S2F, S2G, S2H and 5H, the non-nuclear Z-DNA intensity per cell (cytoplasmic space including cytosol, mitochondria, and other organelles) was calculated from the whole cell area minus the DAPI positive area. Z-DNA signals colocalizing with TOMM20 signals were quantified as mitochondrial Z-DNA, while the remaining non-nuclear Z-DNA staining was quantified as cytosolic Z-DNA. cGAS and Cellular Z-DNA intensity were quantified from Nikon .nd2 files using NIS-Elements AR 5.21.02.

### Flow cytometry

The supernatants from cardiac isolations were centrifuged at 300 x g, 4°C for 5 min with the centrifuge brake off to pellet the non-myocyte cells. The cell pellet was washed with 1 x PBS containing 5% FBS, resuspended in 1 x PBS with Fc Shield (TONBO, 70-0161-M001) and Ghost Dye Red 710 (TONBO, 13-0871-T100), and then loaded at 50 µL/well onto Laminar Wash Plates (Curiox, 96-DC-CL-05). The cells were left on ice for 25 min in dark to settle and then washed using Curiox Laminar Wash HT2000 system at the flow rate of 5 µL/s for 7 times. After wash, the cells were left in the wells with 25 µL of wash buffer. Antibodies were diluted in 1 x PBS containing 5% FBS as 2 x working solution, and 25 µL of the 2 x antibodies were loaded into each well. The cells were incubated with antibodies on ice for 30 min in dark, and then wash as previously described for 12 times. After wash, the cells were collected in polystyrene round-bottom 12 x 75 mm tubes for flow cytometry analysis on Cytek^®^ Aurora 5-Laser Spectral Cytometer.

### Echocardiography

Mice were depilated at the chest area the day before echocardiography measurements. Anesthesia was induced by placing mice in a box with inhaled isoflurane at 2-3%. Afterwards, light anesthesia was maintained using a nose cone with inhaled isoflurane at 0.5-1%. Mice were immobilized on a heated stage, and continuous electric cardiogram (ECG), respiration, and temperature (via rectal probe) were monitored. Light anesthesia and core temperature were maintained to ensure near physiological status with heart rate at range of 470-520 beats per minute. Transthoracic echocardiography was performed using a VisualSonics Vevo 3100 system with a MX550D imaging transducer (center frequency of 40MHz). Parasternal long-axis (B-mode) and parasternal short-axis (M-mode) views of each animal were taken. Vevo LAB cardiac analysis package was used to analyze the data.

### Histology

Mice were euthanized and tissues were washed in PBS and incubated for 24 h in 10% formalin and then transferred to 70% ethanol. Tissue embedding, sectioning, and staining were performed at AML Laboratories in St. Augustine, FL. Images of the H&E staining and Picrosirius red (P. Red) slides were acquired on ECLIPSE Ts2 (Nikon) with a 20x or 40x objective and NIS-Elements D 5.02.01 (Nikon) software. Blinded scoring of 9 independent fields of view from 6 hearts per genotype and treatment was performed to determine percent of cardiomyocytes exhibiting vacuolization post-Doxo. Collagen area was quantified using ImageJ.

### Immunofluorescence staining in cardiac sections

Formalin-Fixed Paraffin-Embedded (FFPE) tissue sections were baked for 30 min, deparaffinized in xylene, and rehydrated by serial passage through graded concentrations of ethanol. Endogenous peroxidase in tissues was blocked with 0.9% H_2_O_2_/methanol for 40 min. Multiple (5) sequential heat-induced epitope retrieval (HIER) treatments were performed for 20 min each at 92-96°C in Citrate pH 6.0 buffer. After each HIER cycle in Citrate pH 6.0 buffer, sections were rinsed with diH_2_O and cooled at RT for 20 min. Tissue sections were circled with hydrophobic barrier pens and blocked with 20% normal goat serum for 20 min before incubation with primary antibody for 20 min. Then, sections were rinsed with TBST for 10 min and incubated with HRP-conjugated secondary (A120-501P) for 20 min, followed by another 10 min rinse in TBST. Incubation with Opal fluorophore Opal690 was then done for 10 min, followed by a 10 min rinse in diH_2_O. Bound primary and secondary antibodies were then subject to the next HIER treatment (as aforementioned) for 20 min. After staining with the Opal fluorophore, tissue specimens were stained with DAPI for 10 min and mounted in VECTASHIELD Vibrance Antifade Mounting Medium (ThermoFisher Scientific). Vectra Polaris Automated Quantitative Pathology Imaging System (Akoya Biosciences) was used for multispectral imaging at 40× magnification. Qupath^122^ and ImageJ were used to export the images and to quantify the ATP5A1 fluorescence intensity.

### RNA sequencing and bioinformatic analyses

Total cellular RNA from WT, *Tfam^+/-^*, *Zbp1^-/-^* and *Tfam^+/-^Zbp1^-/-^* MEFs was prepared using Quick-RNA microprep kit (Zymo Research, R1051) and total cellular RNA from cardiomyocytes isolated from vehicle-and Doxo-treated mice was prepared using Direct-zol RNA miniprep kit (Zymo Research, R2052). Next-generation RNA sequencing procedure for both sets of samples were performed at Texas A&M University Bioinformatics Core. RNA sequencing data were analyzed using BaseSpace Sequence Hub (Illmina). In brief, STAR algorithm of RNAseq Alignment V2.0.0 software was utilized to align the results to reference genome Mus musculus/mm10 (RefSeq), then RNA-seq Differential Expression V1.0.0 software was used to obtain raw gene expression files and determine statistically significant changes in gene expression in *Tfam^+/-^*, *Zbp1^-/-^* and *Tfam^+/-^Zbp1^-/-^* MEFs relative to WT, or cardiomyocytes from Doxo-treated mice related to vehicle-treated mice. Heat maps were generated using GraphPad Prism. For Figures 5A and S5A, GSE40289 and GSE79413 datasets from the Gene Expression Omnibus (GEO) were analyzed using Gene Set Enrichment Analysis software (GSEA, Broad Institute) to identify differentially regulated pathways, Set Comparison Appyter^123^ and Enrichr^124–126^ to identify the interactions between datasets and overlapping genes. For Figure 6B and 6D, RNA-seq data from cardiomyocytes isolated from adult mouse hearts post Doxo challenge compared to vehicle controls were analyzed using ShinyGo^127^ to generate the dot plots and GSEA to generate the enrichment plots.

## QUANTIFICATION AND STATISTICAL ANALYSIS

Error bars displayed throughout the manuscript represent standard error of the mean (s.e.m.) and were calculated from triplicates unless otherwise indicated. For in vivo experiments, error bars were calculated from the average of triplicate technical replicates of at least 5 animals per point. For microscopy quantification, images were taken throughout the slide of each sample using the DAPI channel to avoid bias of selection. To reduce potential experimental bias, samples for cardiac function (EF, FS, IVS;s, LVID;s, LVPW;s, LVPW;d and LV mass) and histological analyses were blinded to researchers when performing analysis. The identities were only revealed at the final data analysis stage. No randomization or blinding was used for all other animal studies. No statistical method was used to predetermine sample size. Data shown are representative of at least 3 independent experiments, including microscopy images, western blots, gene expression analyses and all other cellular assays. All calculations of significance were determined using GraphPad Prism 9.5.0 software. The significance was determined by a p value of < 0.05, and annotated as *P < 0.05, **P < 0.01, and ***P < 0.001.

## SUPPLEMENTAL TABLES

**Table S1. Normalized counts of differentially expressed genes (p < 0.05) in WT, *Tfam^+/-^*, *Zbp1^-/-^*and *Tfam^+/-^Zbp1^-/-^* MEFs, related to Figure 1**.

**Table S2. Normalized counts of differentially expressed genes (p < 0.05) in cardiomyocytes isolated from vehicle and Doxorubicin challenged mice, related to Figure 6**.

